# Exposure and resistance to lantibiotics impact microbiota composition and function

**DOI:** 10.1101/2023.12.30.573728

**Authors:** Zhenrun J. Zhang, Cody Cole, Huaiying Lin, Chunyu Wu, Fidel Haro, Emma McSpadden, Wilfred A van der Donk, Eric G. Pamer

## Abstract

The intestinal microbiota is composed of hundreds of distinct microbial species that interact with each other and their mammalian host. Antibiotic exposure dramatically impacts microbiota compositions and leads to acquisition of antibiotic-resistance genes. Lantibiotics are ribosomally synthesized and post-translationally modified peptides produced by some bacterial strains to inhibit the growth of competing bacteria. Nisin A is a lantibiotic produced by *Lactococcus lactis* that is commonly added to food products to reduce contamination with Gram-positive pathogens. Little is known, however, about lantibiotic-resistance of commensal bacteria inhabiting the human intestine. Herein, we demonstrate that Nisin A administration to mice alters fecal microbiome compositions and the concentration of taurine-conjugated primary bile acids. Lantibiotic Resistance System genes (LRS) are encoded by lantibiotic-producing bacterial strains but, we show, are also prevalent in microbiomes across human cohorts spanning vastly different lifestyles and 5 continents. Bacterial strains encoding LRS have enhanced *in vivo* fitness upon dietary exposure to Nisin A but reduced fitness in the absence of lantibiotic pressure. Differential binding of host derived, secreted IgA contributes to fitness discordance between bacterial strains encoding or lacking LRS. Although LRS are associated with mobile genetic elements, sequence comparisons of LRS encoded by distinct bacterial species suggest they have been long-term components of their respective genomes. Our study reveals the prevalence, abundance and physiologic significance of an underappreciated subset of antimicrobial resistance genes encoded by commensal bacterial species constituting the human gut microbiome, and provides insights that will guide development of microbiome augmenting strategies.

## Main

Antimicrobial resistance (AMR) has emerged as a major public health challenge that is limiting treatment and prevention of an increasing range of bacterial infections. Multi-drug resistant (MDR) bacterial pathogens are causing an estimated 2.8 million infections per year in the United States alone, leading to roughly 35,000 deaths and 55 billion healthcare cost each year ^1^. The increasingly common administration of antibiotics and antimicrobials has contributed to the rising prevalence of MDR pathogens. Antibiotic administration to livestock and poultry has also been linked to emergence of MDR pathogens around the globe ^2–4^.

The human gut microbiota contains trillions of bacteria that contribute to defense against food- borne pathogens by providing colonization resistance. Mechanisms of colonization resistance include nutrient competition, direct antagonism, pH and oxygen manipulation, production and conversion of secondary metabolites, stimulation of immune defense, among others ^5–9^. Reduced microbiota density and diversity following antibiotic use increases susceptibility to infections, especially those caused by MDR pathogens ^10–13^. In humans, fecal microbiota transplantation from donors with an intact microbiota or transfer of purified fecal spores from healthy donors can enhance resistance to *Clostridoides difficile* infection ^14,15^. In mice, administration of defined bacterial consortia can reduce susceptibility to enteric infections by bacterial pathogens ^16–18^. For example, a four-bacterial strain consortium prevents and clears Vancomycin Resistant Enterococcus (VRE) by *in vivo* production of the lantibiotic Blauticin ^18–20^.

Lantibiotics are a class of bacterial ribosomally synthesized and post-translationally modified peptides (RiPPs) that have potent antimicrobial properties and have been widely used in the food industry. Nisin, the prototypical lantibiotic derived from *Lactococcus lactis*, has been used as a food preservative for decades and is generally regarded as safe (GRAS) ^21,22^. The impact of Nisin on the human gut microbiome, however, remains largely undefined. Nisin-like lantibiotics bind to lipid II, a key precursor for bacterial cell wall synthesis, and form pores in the bacterial membrane ^23–25^. Lantibiotic-producing bacteria encode resistance genes, including the ABC transporter *lanFEG*, the lantibiotic immunity protein *lanI*, and nisin resistance protein NSR ^26^. However, the prevalence of lantibiotic resistance system genes (LRS) in the human gut microbiome and whether LRS impact host and microbiota physiology remains unknown.

We demonstrate that dietary Nisin, at concentrations commonly encountered by humans ingesting processed foods, altered the murine gut microbiome and metabolome, diminishing subsets of bacterial taxa and increasing the concentration of taurine-conjugated primary bile acids. LRS are abundant and prevalent in human gut microbiomes across various human cohorts. LRS associated with a two-component system, recombinases and peroxide stress response genes in bacterial genomes, but they co-evolve with the bacterial host and their transcription is constitutive and not responsive to exogenous lantibiotic administration. LRS provides resistance against diet- and microbiota-derived lantibiotics *in vitro* and *in vivo*, and against host-derived Antimicrobial Peptides (AMPs) *in vitro*. However, in the absence of lantibiotic pressure, bacterial strains encoding LRS have reduced *in vivo* fitness. Differential binding of host-derived, secreted IgA contributes to fitness discordance between LRS-encoding and LRS-lacking bacteria. Our results suggest that exposure to lantibiotics and, potentially also to AMPs, has driven the prevalence of LRS in commensal bacterial strains belonging to the Bacillota phylum and impacted microbiota compositions and functions.

### Dietary lantibiotic impacts gut microbiome and metabolome

Nisin, the prototypical lantibiotic, is widely used as food preservative for processed food for decades ^21,22^. While it is generally regarded as safe, Nisin is an antimicrobial peptide, and how it might impact gut microbiome remains understudied. We showed previously that Nisin-like lantibiotics impacted differentially to various gut microbes ^19,20^, and we set out to characterize how dietary Nisin might impact the composition and function of gut microbiome *in vivo*. We supplemented the drinking water of Specific-Pathogen Free (SPF) mice with 10 mg/L Nisin, a concentration commonly found in processed food and drinks ^27^, while the other group of SPF mice were maintained with plain water as control. Fecal samples of both groups were collected over 14 days and samples were processed for metagenomic sequencing and analyzed with MetaPhlAn4 ^28^. We found that Nisin supplementation did not shift overall structure of the gut microbiome on Principle Coordinate Analysis (PCoA) plot with Bray-Curtis dissimilarity metric (Extended Data Fig. 1a), and overall alpha diversity among groups were not significantly changed (Extended Data Fig. 1b,c,d). However, LEfSe analysis found that certain bacterial taxa were differentially diminished upon dietary Nisin supplementation, either comparing Nisin supplementation group versus Control group (Fig. 1a) or comparing Day 14 versus Day 0 in Nisin supplementation group (Fig. 1b). Specifically, compared to Control group, Nisin supplementation diminished the abundance of *Turicibacter* genus and species, an *Acutalibacter* species, as well as a few unnamed genera and species in Oscillospiraceae (Fig. 1a,b); on the other hand, Nisin supplementation consistently enriched for *Lactobacillus* genus, especially *Lactobacillus johnsonii* in both comparison (Fig. 1a,b). This suggests that dietary Nisin differentially impacted the relative abundance of certain Bacillota taxa in gut microbiome.

**Figure 1.**
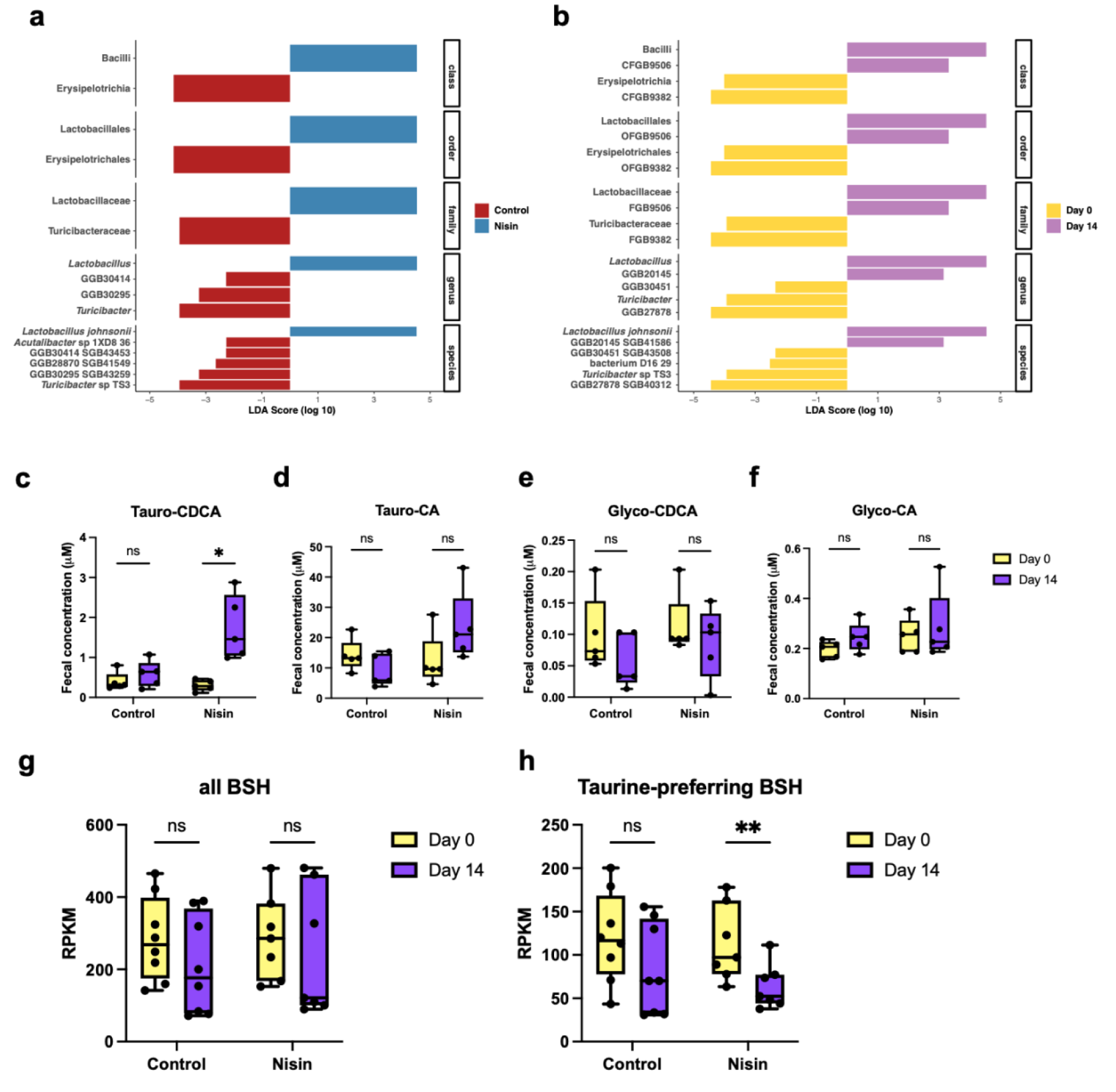
Dietary lantibiotic impacts gut microbiome and metabolome. **(a,b)** LEfSe analysis comparing fecal metagenomes of SPF mice supplemented with 10 mg/L Nisin versus controls (a) or comparing fecal metagenomes of SPF mice 14 days after supplemented with 10 mg/L Nisin versus those before Nisin supplementation (b). Taxa significantly enriched in each group (adjusted *P* value < 0.05) were shown faceted by taxon levels. **(c,d,e,f)** Fecal concentration measurement of tauro-chenodeoxycholic acid (c), tauro-cholic acid (d), glycol-chenodeoxycholic acid (e), and glycol-cholic acid (f) from SPF mice supplemented without (Control) or with 10 mg/L Nisin before (Day 0) or after 14 days of supplementation (Day 14). **(g,h)** RPKM of metagenomic reads mapped to all bile salt hydrolase genes (g) or to taurine-preferring bile salt hydrolase genes (h) from SPF mice supplemented without (Control) or with 10 mg/L Nisin before (Day 0) or after 14 days of supplementation (Day 14). Paired t-tests with Holm-Šídák method for **c-h**. \**P* < 0.05, \*\**P* < 0.01, \*\*\**P* < 0.001; ns, not significant (*P* > 0.05). For **c-h**, box edges indicate 25 and 75 percentiles, horizontal line indicates median, and whiskers indicate min and max.

Gut resident Gram-positive bacteria, which are sensitive to lantibiotics due to lack of outer membrane, are prominent producers of short-chain fatty acids (SCFAs) ^29,30^ and converters of secondary bile acids ^31–33^. To measure the impact of dietary Nisin on biochemical function of the gut microbiome, we also performed metabolomic characterization on Nisin-exposed fecal samples as well as controls. While the concentrations of SCFAs were not significantly changed by Nisin treatment (Extended Data Fig. 1e,f,g), profiles of bile acids were altered. Of note, taurine-conjugated primary bile acids, especially taurochenodeoxycholic acid, but not glycine-conjugated primary bile acids, increased upon Nisin supplementation (Fig. 1c,d,e,f). We hypothesized that taurine-preferring bile salt hydrolases (BSH) ^34^ encoded in the gut metagenome were diminished upon exposure to dietary Nisin. To this end, we performed metagenomic mapping of taurine-preferring BSH genes and all BSH genes in both groups. We found that the relative abundance of taurine-preferring BSH decreased in Nisin supplementation group, corroborating our metabolomic observation, while the relative abundance of all BSH did not change significantly (Fig. 1g,h). In summary, dietary Nisin shifted the composition of microbiome and metabolomic profile in mice gut.

### Lantibiotic resistance systems are prevalent among human gut bacteria

To further investigate how lantibiotics might impact human gut bacteria, we focused on mining for lantibiotic resistance systems (LRS) as well as lantibiotic biosynthetic gene clusters (BGC) in human gut bacterial genomes. Duchossois Family Institute symbiotic bacteria collection (DFI collection) has more than 1,500 bacterial isolates from healthy human donors that are purified and whole-genome sequenced. For 1,576 isolates surveyed, we found 716 unique strains of bacteria according to their average nucleotide identity (ANI) to other isolates within the same species (ANI cutoff at more than 99.9%). In search for the presence of genes within Nisin-like lantibiotic biosynthetic gene cluster (*lanABCFEGI*) among their genomes, we found wide-spread presence of LRS genes (*lanFEG* and *lanI*) among these bacteria (247/716, 34.5%) (Figure 2a; Extended Data Table 1), especially in the Bacillota (previously Firmicutes) phylum, and 99 of them contain both *lanFEG* and *lanI* in their genomes. In stark contrast, lantibiotic structural and biosynthetic genes *lanA* and *lanBC* were rare among human gut bacteria (18/716, 2.5%), and only 7 of them had presence of both *lanA* and *lanBC* in their genome (Figure 2a; Extended Data Table 1). To exclude potential bias caused by culturing approach that establishes the DFI collection, we performed metagenomic sequencing on all the samples from healthy human donors and mapped the reads to lantibiotic structural gene *lanA* and LRS genes (Extended Data File 1-5). Consistently, we observed that LRS genes are present among all donor samples, and their relative abundances are much higher than that of *lanA*. In fact, we could detect reads mapped to *lanA* in only about half of our donor cohort (Extended Data Figure S2A). To further extend our observation, we performed similar analyses on publicly available gut metagenomes of different healthy human cohorts, including Human Microbiome Project (HMP) ^35^, Metagenomics of the Human Intestinal Tract (MetaHIT) ^36^, a Japanese cohort ^37^, a Hadza cohort ^38^ and a Yanomami cohort ^39^, altogether spanning five continents and diverse lifestyles. Again, we found that LRS genes are present and more abundant than lantibiotic structural gene *lanA* across all cohorts, while *lanA* gene was detected in only one sample within the Hadza cohort and is absent in Japanese and Yanomami cohorts (Figure 2b). All the observations above suggest that human gut bacteria harbor abundant LRS genes whereas lantibiotic structural genes required to produce active lantibiotics are much less abundant, even rarely present.

**Figure 2.**
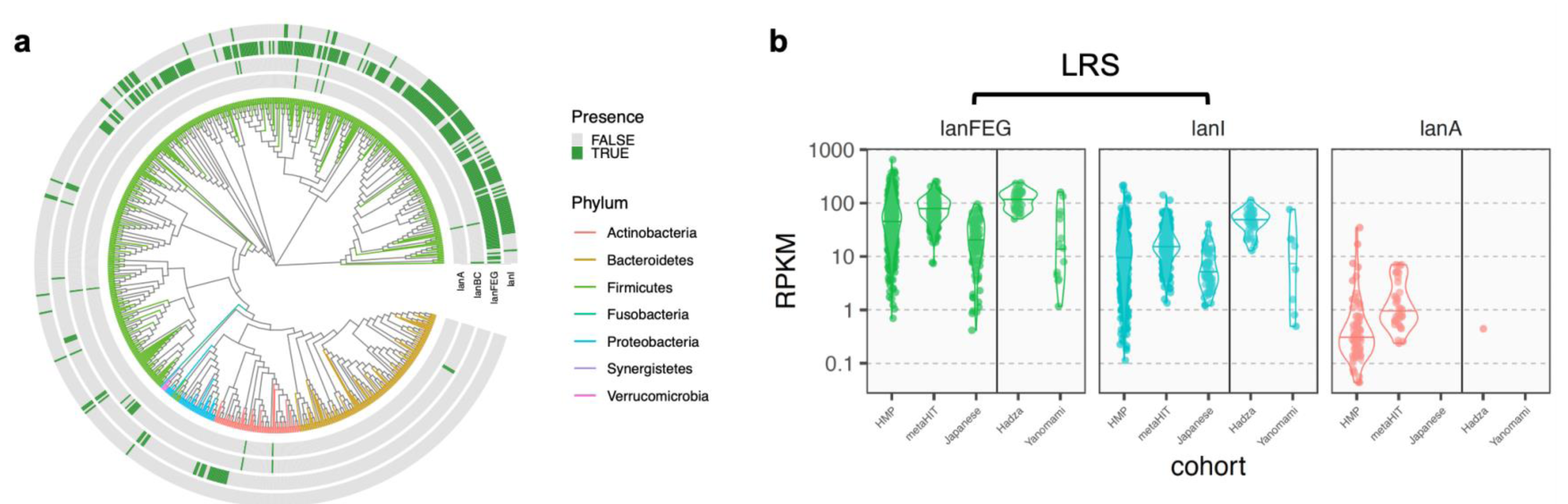
Lantibiotic resistance systems (LRS) are prevalent among human gut bacteria. **(a)** Distribution of lantibiotic structural gene *lanA*, biosynthetic genes *lanBC*, LRS genes *lanFEG* and *lanI* in unique isolates of DFI collection. Isolates were organized in a phylogenetic tree based on their 16S rRNA sequences, colored by their phylum identity. Color of boxes in outer rings indicate presence (green) or absence (gray) of respective genes. **(b)** RPKM of metagenomic reads from Human Microbiome Project (HMP), Metagenomics of the Human Intestinal Tract (MetaHIT), a Japanese cohort, a Hadza cohort and a Yanomami cohort mapped to *lanFEG*, *lanI* and *lanA*, respectively. Vertical line in each graph separates industrialized lifestyle (left) and hunter-gatherer lifestyle (right).

### LRS are associated with two-component system and recombinase

We next set out to investigate the genomic context within which LRS genes occur. Within the isolates that contain LRS in their genomes, we observed a consistent pattern in the organization of LRS genes. For isolates that have only one LRS locus, the three-component ABC transporter system *lanFEG* are organized in tandem and in that order, and the lantibiotic immunity protein gene *lanI* is directly downstream of *lanFEG* if *lanI* is present. A two-component regulatory system *lanRK* is always present downstream of LRS genes, presumably regulating the expression of LRS (Figure 3). We found more than one LRS locus in the genomes of 34 unique isolates in DFI collection, each organized as described (*lanFEG(I)RK*), except for *Blautia wexlerae* isolates, which have a *lanFEG*-only operon upstream of a complete LRS loci, separated by one open reading frame (ORF) (Figure 3). These two copies of *lanFEG* are not identical, sharing only about 70% identity. We have also observed some pseudogenization events within LRS, including premature stop codon in *lanI* of *Blautia schinkii* MSK.15.25 and in *lanR* of *Blautia massiliensis* MSK.18.38, breaking up the genes into two ORFs (Figure 3).

**Figure 3.**
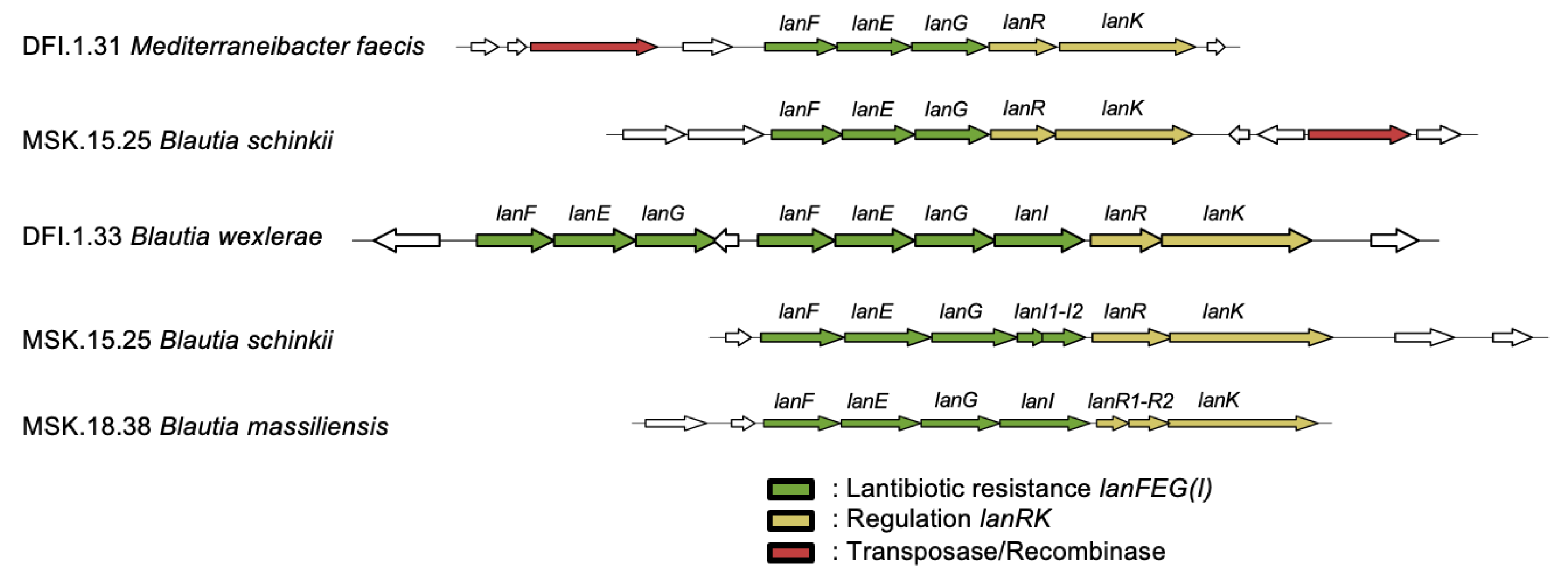
LRS are associated with two-component system and recombinase. Representative LRS operons from isolates in DFI collection are shown. Identities of bacterial genomes are labeled on the left. LRS genes and associated two-component system genes are annotated on top. LRS, two- component system genes, and transposase/recombinase are colored.

To further investigate the genomic context of LRS operons, we examined genes in the flanking regions of LRS loci. We numerated gene annotations of 10 ORFs flanking all LRS loci; besides unannotated hypothetical proteins, we frequently found the presence of transposases or recombinases (Figures 3 and Extended Data Figure 3a). The most frequent annotated gene in the LRS flanking regions is transposase from transposon Tn916 (Extended Data Figure 3a), occurring in about 20% of LRS loci. Tyrosine recombinase XerC also ranked among the top 15 frequent genes flanking LRS (Extended Data Figure 3a). Interestingly, we also found frequent presence of genes related to peroxide stress resistance, including putative peroxiredoxin bcp, peroxide stress resistance protein YaaA, and peroxide-responsive repressor PerR (Extended Data Figure 3a). The fact that LRS loci are frequently flanked by transposases or recombinases suggest LRS may undergo horizontal gene transfer among human gut bacteria. To test this hypothesis, we built distance matrices using sequences of full-length 16S rRNA and each of the LRS genes among the LRS-containing isolates in DFI collection, and then assessed the likelihood of co-speciation between 16S rRNA and LRS genes by comparing the similarity of these matrices. Procrustes analyses after Principle Coordinate Analysis superimposition ^40^ found that the co-speciation of LRS genes with microbes were statistically significant (Extended Data Figure 3b,c,d,e), suggesting that LRS genes are fixated on the genomes of human gut microbes, even though they may have horizontally transferred among gut commensals on an evolutionary timescale.

### LRS provides resistance against lantibiotics and antimicrobial peptides in vitro

To investigate whether LRS discovered *in silico* have physiological significance, we set out to test and compare sensitivity of human gut bacterial isolates towards lantibiotics. We selected pairs of bacterial isolates of the same species that are discordant in the presence of LRS in their genomes for comparison. These isolates were cultured in 96-well plates with a concentration gradient of Nisin, a diet-derived lantibiotic, and Blauticin, a gut commensal-derived lantibiotic ^19,20^, both heterologously expressed and purified ^19,41^, and determined their minimal inhibitory concentrations (MIC). We found that isolates with LRS in their genomes have consistent higher resistance against both lantibiotics compared to their LRS-negative counterparts (Figure 4a,b,c, Extended Data Figure 4a,b,c). Moreover, *lanFEG*+ *lanI*+ isolates had increased lantibiotic resistance than *lanFEG*+ *lanI*- counterparts, suggesting that *lanI* gene provides additional resistance to that provided by *lanFEG* (Figure 3c, Extended Data Figure 3c).

**Figure 4.**
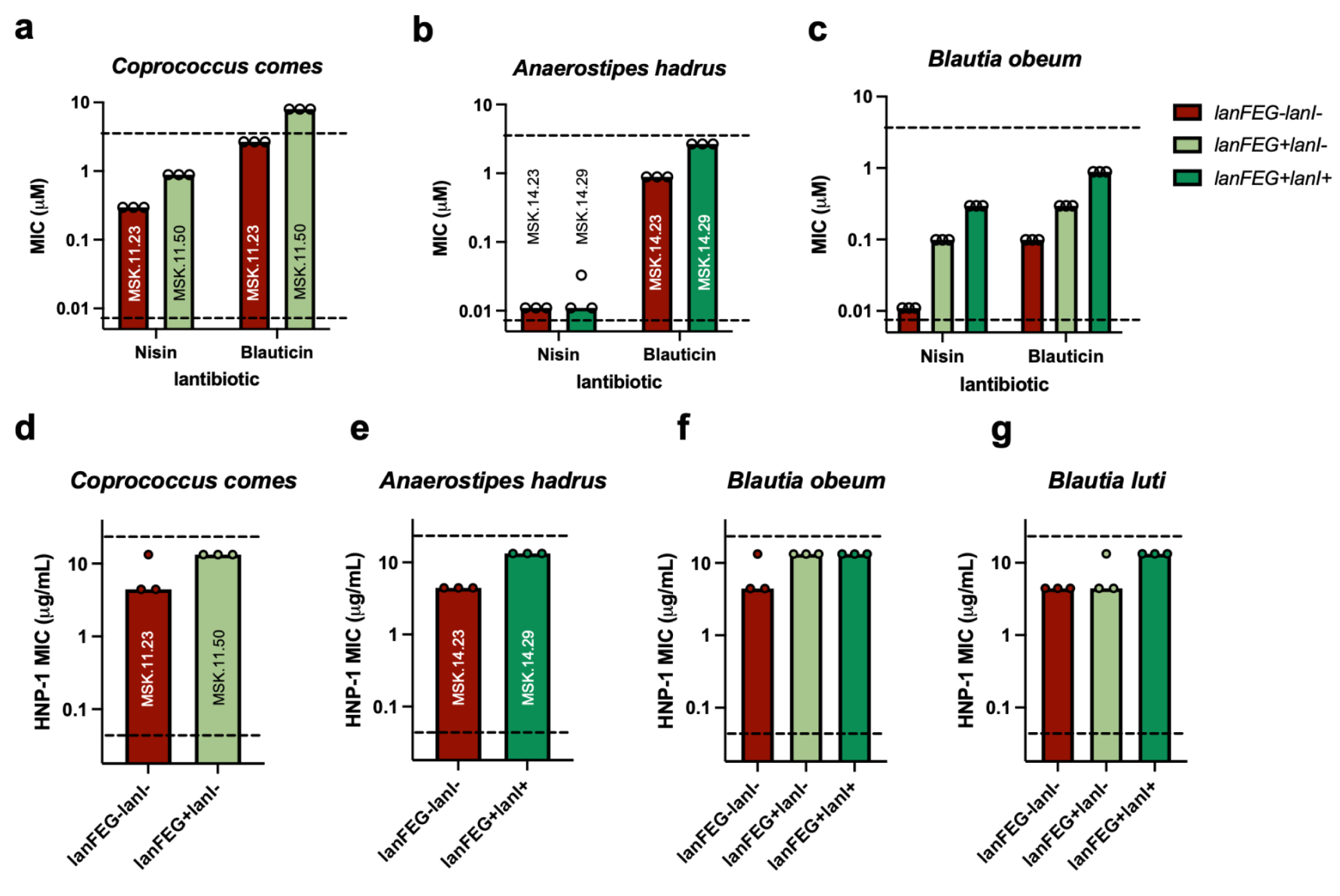
LRS provides resistance against lantibiotics and antimicrobial peptides *in vitro*. **(a,b,c)** Minimal Inhibitory Concentration (MIC) of Nisin and Blauticin on *Coprococcus comes* (a), *Anaerostipes hadrus* (b) and *Blautia obeum* (c) isolates from human gut. Each circle represents one independent measurement. Horizontal dashed lines indicate upper and lower limit of concentration tested, respectively. Color indicates presence or absence of LRS genes. Identities of *C. comes* and *A. hadrus* isolates are annotated. **(d,e,f,g)** Minimal Inhibitory Concentration (MIC) of human α-defensin HNP-1 on *Coprococcus comes* (d), *Anaerostipes hadrus* (e), *Blautia obeum* (f) and *Blautia luti* (g) isolates from human gut. Each circle represents one independent measurement. Horizontal dashed lines indicate upper and lower limit of concentration tested, respectively. Color indicates presence or absence of LRS genes. Identities of *C. comes* and *A. hadrus* isolates are annotated.

Previous research suggests that many lantibiotic BGCs that contain LRS genes are silent under laboratory conditions ^42,43^, while Nisin BGC in *L. lactis* is self-inducible by Nisin ^44–46^. To confirm whether LRS genes are actively expressed and to test whether LRS genes are inducible by lantibiotic *in vitro*, we performed quantitative RT-PCR on LRS-positive bacteria cultured in the absence or presence of Nisin at one-third of respective MIC. We observed that LRS genes were expressed *in vitro* at levels comparable to housekeeping gene *ftsZ* (Extended Figure 5a-k), although their expression levels were not induced by exogenous Nisin (Extended Figure 5l-v). These data suggest that *in silico* identified LRS are expressed and functional *in vitro*.

**Figure 5.**
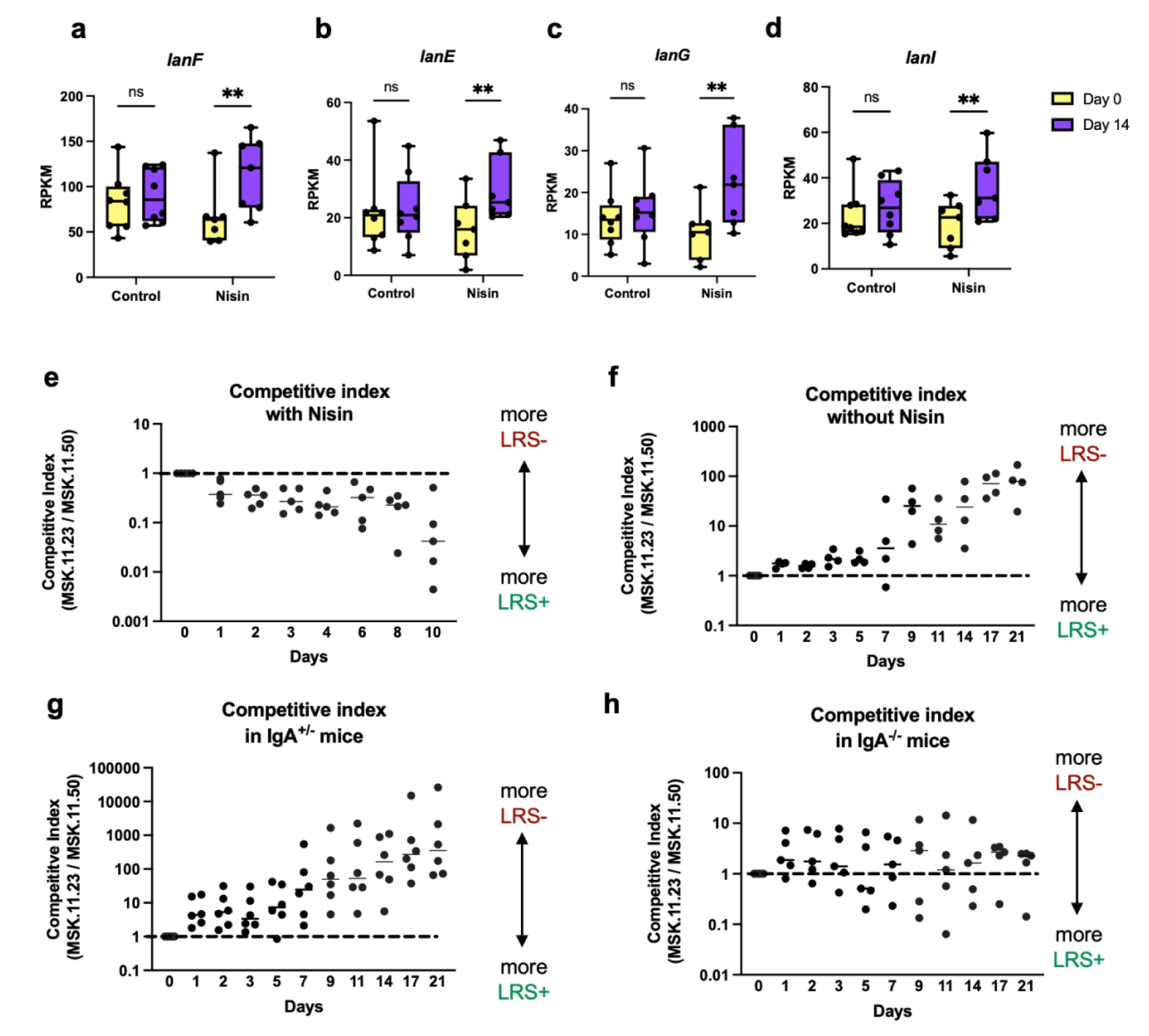
LRS modulate fitness dynamics of human gut commensals *in vivo*. **(a,b,c,d)** RPKM of metagenomic reads mapped to *lanF* (a), *lanE* (b), *lanG* (c) and *lanI* (d), respectively, from SPF mice supplemented without (Control) or with 10 mg/L Nisin before (Day 0) or after 14 days of supplementation (Day 14). **(e,f)** Competitive index of *C. comes* MSK.11.23 (LRS-negative) over MSK.11.50 (LRS-positive) in gnotobiotic mice with (e) or without Nisin supplementation (f), colonized with 1:1 ratio of *C. comes* pair and two Bacteroidetes strains on Day 0. Each point represents one mouse. Dashed line indicates starting competitive index (1). Solid line plots median at each time point. **(g,h)** Competitive index of *C. comes* MSK.11.23 (LRS-negative) over MSK.11.50 (LRS-positive) in IgA^+/-^ (g) or IgA^-/-^ (h) gnotobiotic mice without Nisin supplementation, colonized with 1:1 ratio of *C. comes* pair and two Bacteroidetes strain on Day 0. Each point represents one mouse. Dashed line indicates starting competitive index (1). Solid line plots median at each time point. Paired t-tests with Holm-Šídák method for **a-d**. \**P* < 0.05, \*\**P* < 0.01, \*\*\**P* < 0.001; ns, not significant (*P* > 0.05). For **a-d**, box edges indicate 25 and 75 percentiles, horizontal line indicates median, and whiskers indicate min and max.

Mammalian intestine produces a slew of antimicrobial peptides (AMPs) in the gut to restrict dysbiotic expansion of gut microbes and maintain host-microbe homeostasis ^47–49^. These AMPs include short peptides like α-defensins, β-defensins and LL-37, among many other longer peptides and proteins. α-defensins and β-defensins contain multiple intramolecular disulfide linkages, while LL-37 is a linear helical peptide. We hypothesized that LRS also provides cross-protection against functionally and structurally similar cationic AMPs. We challenged aforementioned pairs of LRS-discordant bacterial isolates with LL-37, human α-defensin HNP-1, human β-defensins hBD-1 and hBD-4, respectively, to measure their MIC. We found that LRS- positive bacteria have consistent higher resistance against HNP-1 (Figure 4d-g), while having same MIC against LL-37 compared to their LRS-negative counterparts (Extended Data Figure 4d- g). None of the bacteria was sensitive to hBD-1 or hBD-4 at highest concentration tested (data not shown). These data suggest that LRS also provide cross-resistance against human α-defensin, further implying their physiological significance in resisting AMPs.

### LRS modulate fitness dynamics of human gut commensals in vivo

To investigate whether resistance provided by LRS *in vitro* translates to physiological significance *in vivo*, we first tested whether LRS among gut bacteria responds to dietary lantibiotic *in vivo*. We found that the relative abundance of LRS genes in the metagenomes of SPF mice supplemented with Nisin were significantly enriched compared to controls after 14 days (Figure 5a-d). This suggests that dietary Nisin shifted the functional profile of gut microbiome by increasing the relative abundance of LRS genes.

An interesting observation in our experiment is that we have isolated pairs of human gut commensals that are discordant in the presence of LRS from the same human donor; for example, *Coprococcus comes* MSK.11.23 and MSK.11.50 (Figure 4a,d) are both isolated from the same human donor FC0011, and *Anaerostipes hadrus* MSK.14.23 and MSK.14.29 (Figure 4b,e) are both isolated from the same human donor FC0014. Given higher resistance to both lantibiotics and host-derived AMPs, the LRS-containing isolate MSK.21.50 would have been expected to outcompete its LRS-lacking counterpart MSK.21.23 *in vivo*. The fact that we have isolated both LRS-positive and LRS-negative strains from the same donors suggest certain mechanisms allow them to co-exist in the same gut community, or counter-select LRS-containing bacteria *in vivo*. Besides, among the HMP cohort and metaHIT cohort that have an adequate number of individuals whose metagenomes contain *lanA* for analysis, the relative abundance of LRS genes were not correlated with that of *lanA*, except for the case of *lanI* in metaHIT cohort, which showed positive correlation with *lanA* (Extended Data Figure 2b,c). This observation further suggests that metagenome-encoded lantibiotics may not increase the relative abundance of LRS genes in human gut metagenomes.

We then set out to test whether LRS modulates bacterial fitness *in vivo*. To avoid potential complications from a complex microbiome, we decided to use gnotobiotic mice model colonized with defined bacterial communities to investigate bacterial fitness dynamics *in vivo*. *Coprococcus comes* could not monocolonize the gut of Germ-Free (GF) mice (data not shown), so we included two mouse-derived Bacteroidota (previously Bacteroidetes) isolates, one *Bacteroides sartorii* and the other *Parabacteroides distasonis*, to form a stable bacterial consortium *in vitro* and *in vivo* ^18,20^. We confirmed that *C. comes* LRS-negative MSK.11.23 and LRS-positive MSK.11.50 had similar growth rates *in vitro* (Extended Data Figure 6a), as well as those between *A. hadrus* LRS- negative MSK.14.23 and LRS-positive MSK.14.29 (Extended Data Figure 6b). The two Bacteroidota isolates did not affect relative competitiveness between the two *C. comes* isolates or the two *A. hadrus* isolates during consecutive culturing and passage for 7 days *in vitro* (Extended Data Figure 6c,d). We then colonized Germ-Free mice with the pair of *C. comes* in 1:1 ratio along with the two Bacteroidota isolates with or without Nisin supplementation in drinking water at a concentration commonly found in human processed food (10 mg/L) ^27^. Consistent with their differences in MIC of Nisin *in vitro*, LRS-positive isolate MSK.11.50 slowly outcompeted LRS- negative isolate MSK.11.23 over time (Figure 5e). Similar phenomenon was observed for the pair of *A. hadrus*, as LRS-positive MSK.14.29 outcompeted LRS-negative isolate MSK.14.23 over time with dietary Nisin supplementation (Extended Data Figure 6e). These suggest that LRS provides fitness advantage to gut bacteria in the presence of dietary lantibiotic *in vivo*. Surprisingly, *C. comes* LRS-negative isolate MSK.11.23 slowly outcompeted LRS-positive isolate MSK.11.50 over time in the absence of Nisin supplementation (Figure 5f). Similarly, *A. hadrus* LRS-negative isolate MSK.11.23 slowly outcompeted LRS-positive isolate MSK.11.50 over time without dietary Nisin supplementation (Extended Data Figure 6f). Because these pairs of isolates did not show fitness difference *in vitro* in the absence of lantibiotics (Extended Data Figure 6a-d), we hypothesized that certain host factors absent *in vitro* contribute to fitness advantage of LRS-negative isolates or fitness disadvantage of LRS-positive isolates *in vivo*.

Human and mice guts respond to antigens from gut commensals and secrete IgA, which is important for the immune homeostasis of the host as well as colonization of gut microbes ^50–53^. LRS proteins are transmembrane or membrane-anchored proteins, so they are potentially accessible antigens for the immune cells and are potentially exposed to secreted IgA from the host. Therefore, we hypothesized that secreted IgA provides fitness discrimination between LRS- positive and LRS-negative bacteria. To test this, we used mice whose first axon of both IgA alleles was deleted, thereby B cells cannot switch to IgA (denoted as IgA^-/-^ mice). To this end, we focused on the *C. comes* pair, so we colonized GF IgA^-/-^ mice with the pair of LRS-discordant *C. comes* in 1:1 ratio along with the two Bacteroidota isolates, while GF IgA-heterozygous (IgA^+/-^) mice littermates, which could still produce and secrete IgA, were colonized with the same 4-bacteria consortium to serve as control. We found that while LRS-discordant pair behaved similarly in IgA^+/-^ mice to wild-type mice in that LRS-negative bacteria had fitness advantage (Figure 5g), the fitness difference *in vivo* between the pair of bacteria disappeared in IgA^-/-^ mice (Figure 5h). To see whether IgA binds differently to these bacteria, we performed magnetic enrichment of IgA- bound bacteria from the cecal samples of these gnotobiotic mice, and quantitative PCR to measure the relative abundance of the pair. Surprisingly, we found that compared to pre-sort samples of IgA^+/-^ mice, IgA-bound samples of IgA^+/-^ mice had significantly enriched LRS-negative MSK.11.23 over LRS-positive MSK.11.50, suggesting that MSK.11.23 was bound with IgA more than MSK.11.50 (Extended Data Figure 6g,i). As a control, similar enrichment did not enrich either of the pair from the cecal contents of IgA^-/-^ mice (Extended Data Figure 6h,i). These data suggest that secreted IgA in the mice gut contribute to fitness disparity between LRS-positive and LRS- negative bacteria. In summary, our data suggest that host factors in the gut, including secreted IgA, contribute to the fitness dynamics of bacteria containing or lacking LRS in their genomes.

## Discussion

Nisin as a bacterially derived antimicrobial peptide has been used as food preservative for decades ^21,22^, yet its exact impact on human gut microbiome has been largely understudied. We supplemented dietary Nisin to mice with intact gut microbiome at a concentration commonly found in processed food ^27^, and observed significant compositional and functional changes in gut metagenome and metabolome. Recent studies have found that *Turicibacter* strains differentially deconjugate host bile acids and alters host serum lipid profiles ^54^, while decreased abundance of *Turicibacter* species is repeatedly associated with obesity ^55^ and inflammatory bowel disease ^56^. The increase in taurine-conjugated primary bile acids and concurrent decrease of the relative abundance of certain *Turicibacter* species in microbiome as well as that of taurine-preferring BSH genes in metagenome suggest that dietary lantibiotic selectively depletes certain bacterial taxa that encodes taurine-preferring BSH and leads to alteration of bile acid profile in the gut.

Further investigation of lantibiotic related genes among human gut bacteria leads us to discover widespread prevalence and high relative abundance of LRS in human gut microbiome, especially within Bacillota phylum. On the contrary, lantibiotic structural and biosynthetic genes are rarely present. Interestingly, Hadza cohort, a group with hunter-gatherer lifestyle, has highest relative abundance of LRS on average among cohorts we investigated, even though lantibiotic structural gene *lanA* is rarely present. We hypothesize that their unique dietary structure ^38^ may contain natural lantibiotics or similar antimicrobial peptides, and genetic differences in this cohort may also leads to higher concentration of host-derived antimicrobial peptides, selecting for bacteria that contain LRS which also provides cross-protection.

LRS genes are organized in an operon together with *lanRK* two component regulatory system. Even though previous studies suggest *lanRK* responds to exogenous lantibiotic and regulates expression of lantibiotic biosynthetic gene cluster in lantibiotic-producing bacteria ^44–46^, we found no evidence of them regulating expression of LRS in LRS-containing, lantibiotic-non-producing human gut bacteria. The fact that we observed instances where *lanR* and LRS gene *lanI* undergo pseudogenization events also suggest that LRS and its associated two component system may pose fitness disadvantage in certain circumstances and therefore have undergone counter- selection process. Besides, as we demonstrated that LRS provides cross-protection against host- derived α-defensin, constitutive expression of LRS rather than regulated expression may provide fitness advantage in the gut. While LRS is frequently associated Integrative-Conjugative Element (ICE) genes, it has co-evolved with the host it resides, suggesting that ICE genes may be remnant of past horizontal gene transfer events that happened in evolutionary timescale. This contrasts with another widely disseminated interbacterial defense systems, which are readily transferrable among Bacteroidales species ^57^. LRS is also frequently associated with peroxide stress response genes. Lantibiotics can inhibit cell wall synthesis and form pores on bacterial membrane, both of which are associated with intracellular ROS stress ^58,59^. Proximity of LRS and peroxide stress response genes may help bacteria to co-regulate these two sets of genes to synergistically respond to exogenous stress caused by lantibiotics and AMPs.

LRS increased bacterial resistance to dietary and gut-derived lantibiotics, yet its relative abundance in human gut metagenome did not positively correlate with that of *lanA*, except in the case of *lanI* in metaHIT cohort. This may be due to two-fold reasons: lantibiotic structural and biosynthetic genes are often silent, as bacteria encoding these genes do not express them; LRS genes have additional function as they provide cross-protection against host-derived AMPs. Its cross-protection has selectivity towards AMPs that is similar structurally to lantibiotics like α- defensin, but not linear peptide LL37. Contrary to silent lantibiotic structural and biosynthetic genes in many bacterial hosts ^42,43^, LRS are expressed constitutively *in vitro* at levels on par with housekeep gene.

Dietary Nisin supplementation selects for LRS-containing bacteria *in vivo*, both in complex SPF microbiome and in defined bacterial consortia. Surprisingly, in the absence of Nisin supplementation, LRS-containing strain were outcompeted by LRS-lacking strain of the same species. Their growth and competitiveness were similar *in vitro*, suggesting that host factor(s) *in vivo* contribute to their fitness differences. We found that secreted IgA in the gut contributed to selecting for LRS-lacking strain, as it binds more to LRS-lacking strain. IgA binding and aggregation of gut bacteria has been shown to facilitate colonization of commensals in the gut ^51^. Detailed mechanisms by which LRS-lacking strain can be bound more by IgA or LRS-containing strain excludes IgA binding requires further investigation.

Our studies demonstrate significant impact on gut microbiome by dietary lantibiotic, and further investigation on structural-activity relationship of lantibiotics should help design novel lantibiotics that have improved selectivity towards pathogens but spares normal gut residents ^19^. While worldwide healthcare system is encountering increasing burden of AMR, we have revealed prevalence, abundance and physiological significance of previously underappreciated AMR genes in human gut microbiome, which will help development of novel therapeutics and strategy to cope with AMR ^60–62^.

## Author Contribution

The study was conceived and designed by Z.J.Z. and E.G.P. Z.J.Z. performed all experiments and bioinformatic analyses unless stated otherwise. Z.J.Z. and C.C. performed experiments and bioinformatic analysis of dietary Nisin on SPF mice metagenome and metabolome. H.L. and Z.J.Z. performed bioinformatic analysis on relative abundance of lantibiotic-related genes on human cohorts. Z.J.Z., F.H. and E.M. performed experiments involving Germ-Free mice. C.W. expressed and purified Nisin and Blauticin for *in vitro* assays. The manuscript was written by Z.J.Z. All authors contributed to editing the manuscript and support the conclusions.

## Competing Interests

E.G.P. serves on the advisory board of Diversigen; is an inventor on patent applications WPO2015179437A1, titled “Methods and compositions for reducing Clostridium difficile infection,” and WO2017091753A1, titled “Methods and compositions for reducing vancomycin resistant enterococci infection or colonization”; and receives royalties from Seres Therapeutics, Inc. The other authors are not aware of any affiliations, memberships, funding, or financial holdings that might be perceived as affecting the objectivity of this manuscript.

## Acknowledgments

The authors are grateful to Host-Microbe Metabolomics Facility of Duchossois Family Institute (DFI) for metabolomic analysis, Microbiome Metegenomics Facility of DFI for metagenomic and bacterial whole-genome sequencing, the Bioinformatics Core of DFI for bacterial genome assembly and annotation, and the Symbiotic Strain Bank of DFI for bacterial strain isolation and maintenance. We thank Q. Dong, R. Oliveira, M. Sorbara, E. Littmann, R. Pope and S. Son for helpful discussion. We thank Z. Earley and A. Bendelac for providing IgA^+/-^ and IgA^-/-^ mice. Z.J.Z was supported by The GI Research Foundation early career grant. This work was supported by US National Institutes of Health grants R01 AI095706, P01 CA023766, U01 AI124275, and R01 AI042135 to E.G.P., and by the Duchossois Family Institute of the University of Chicago.

---

**Extended Data Figure 1.**
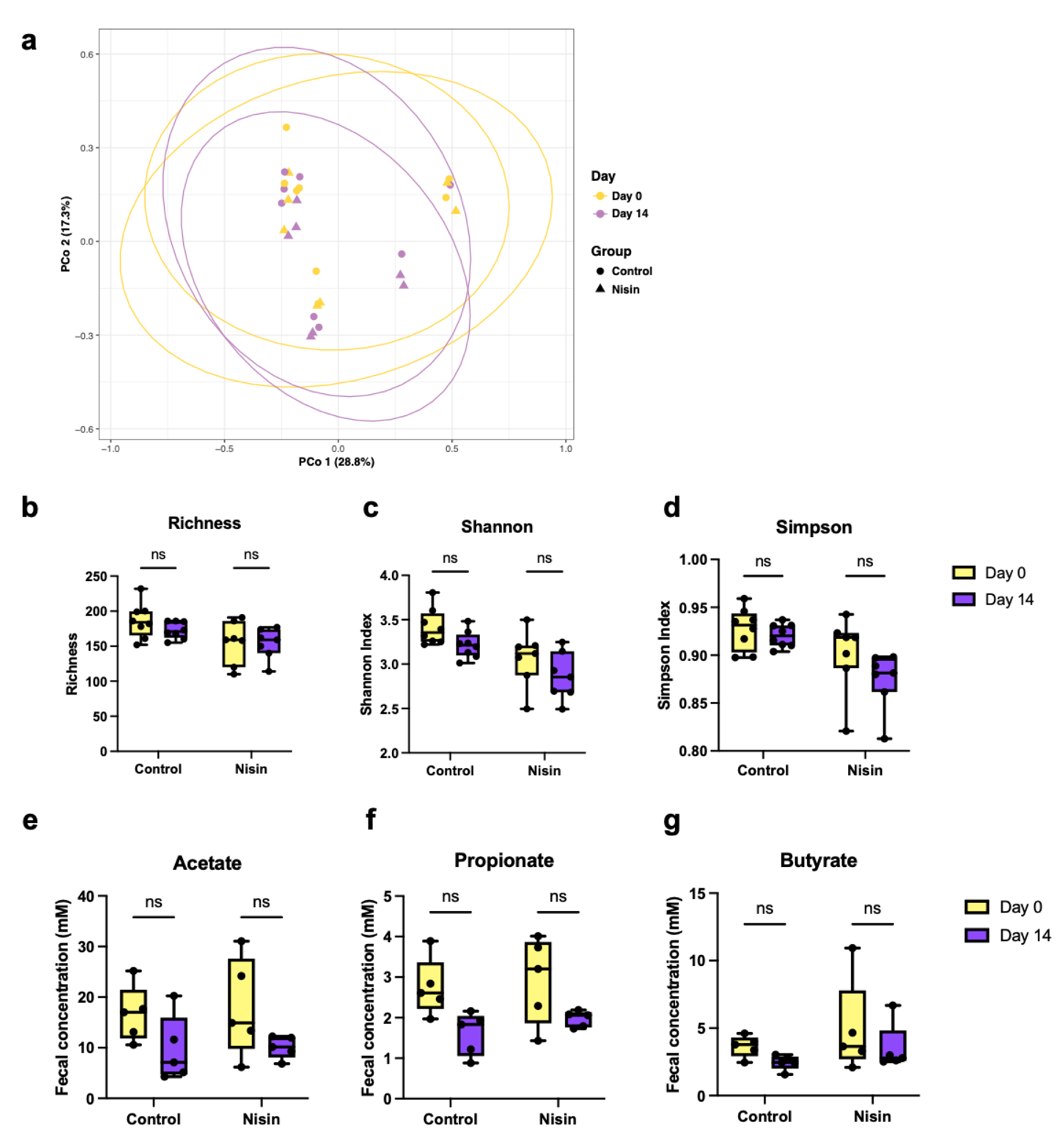
Impact of dietary Nisin on gut microbiome and metabolome. **(a)** PCoA plot of fecal metagenomes of SPF mice without (Control) or with supplemented with 10 mg/L Nisin before (Day 0) or after 14 days of Nisin supplementation (Day 14). Colors indicate supplementation day and shapes indicates supplementation group. Ellipses indicate 95% confidence level of each group. **(b,c,d)** Alpha diversity of fecal metagenomes of SPF mice without (Control) or with supplemented with 10 mg/L Nisin before (Day 0) or after 14 days of Nisin supplementation (Day 14), measured as Richness (b), Shannon Diversity Index (c), or Simpson Diversity Index (d). **(e,f,g)** Fecal concentration measurement of acetate (e), propionate (e), and butyrate (g) from SPF mice supplemented without (Control) or with 10 mg/L Nisin before (Day 0) or after 14 days of supplementation (Day 14). Paired t-tests with Holm-Šídák method for **b-g**. \**P* < 0.05, \*\**P* < 0.01, \*\*\**P* < 0.001; ns, not significant (*P* > 0.05). For **b-g**, box edges indicate 25 and 75 percentiles, horizontal line indicates median, and whiskers indicate min and max.

**Extended Data Figure 2.**
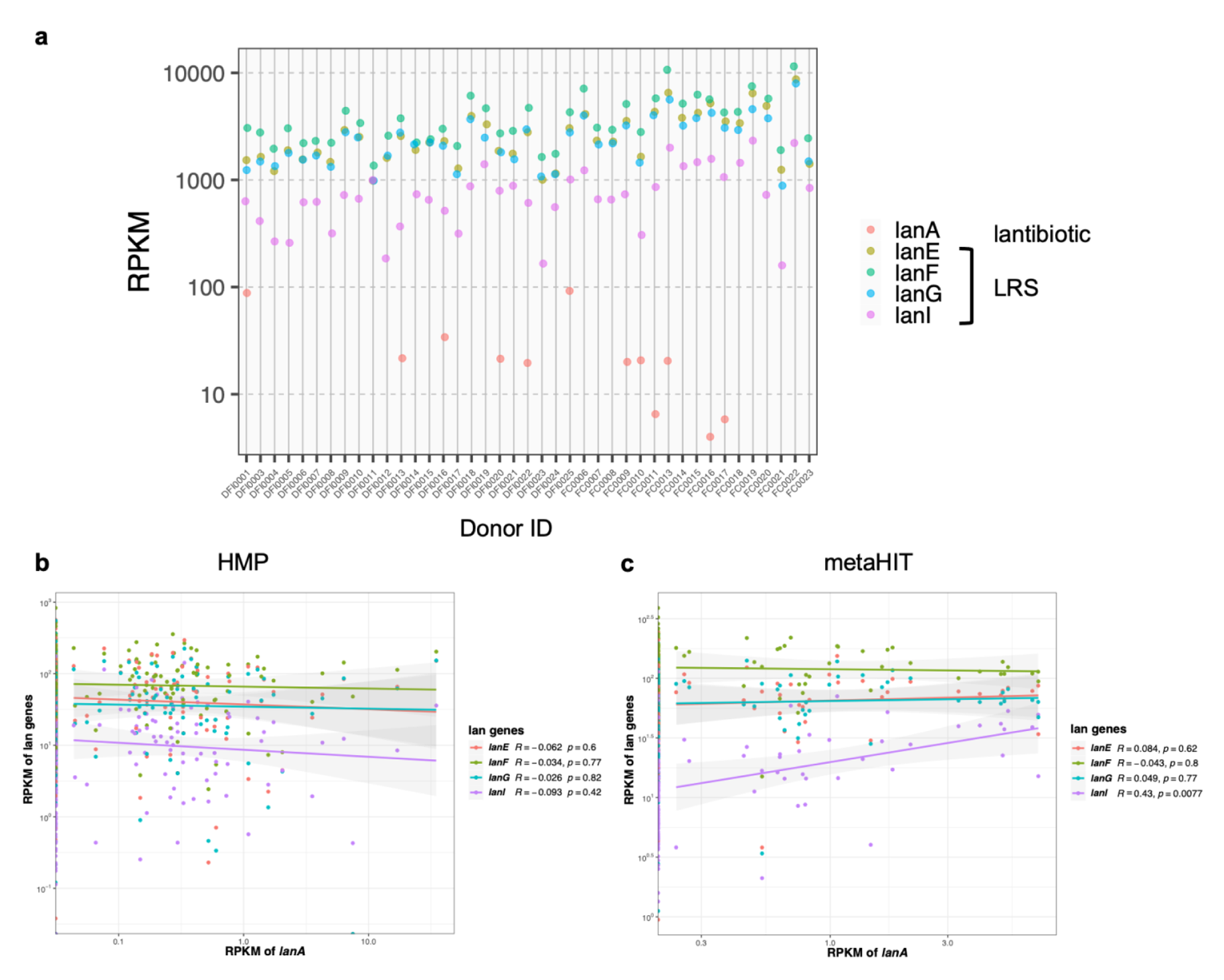
Prevalence of Lantibiotic resistance systems (LRS) among human gut metagenomes. **(a)** RPKM of each LRS gene and *lanA* in metagenomes of each healthy human donor of DFI collection. Each column represents one donor. Colors indicate gene identity. **(b,c)** Correlation between RPKM of each LRS gene and RPKM of *lanA* in HMP cohort (b) or metaHIT cohort (c). Correlation coefficient *R* and significance *p* of each correlation are labeled on the right. Colored lines plot linear regression.

**Extended Data Figure 3.**
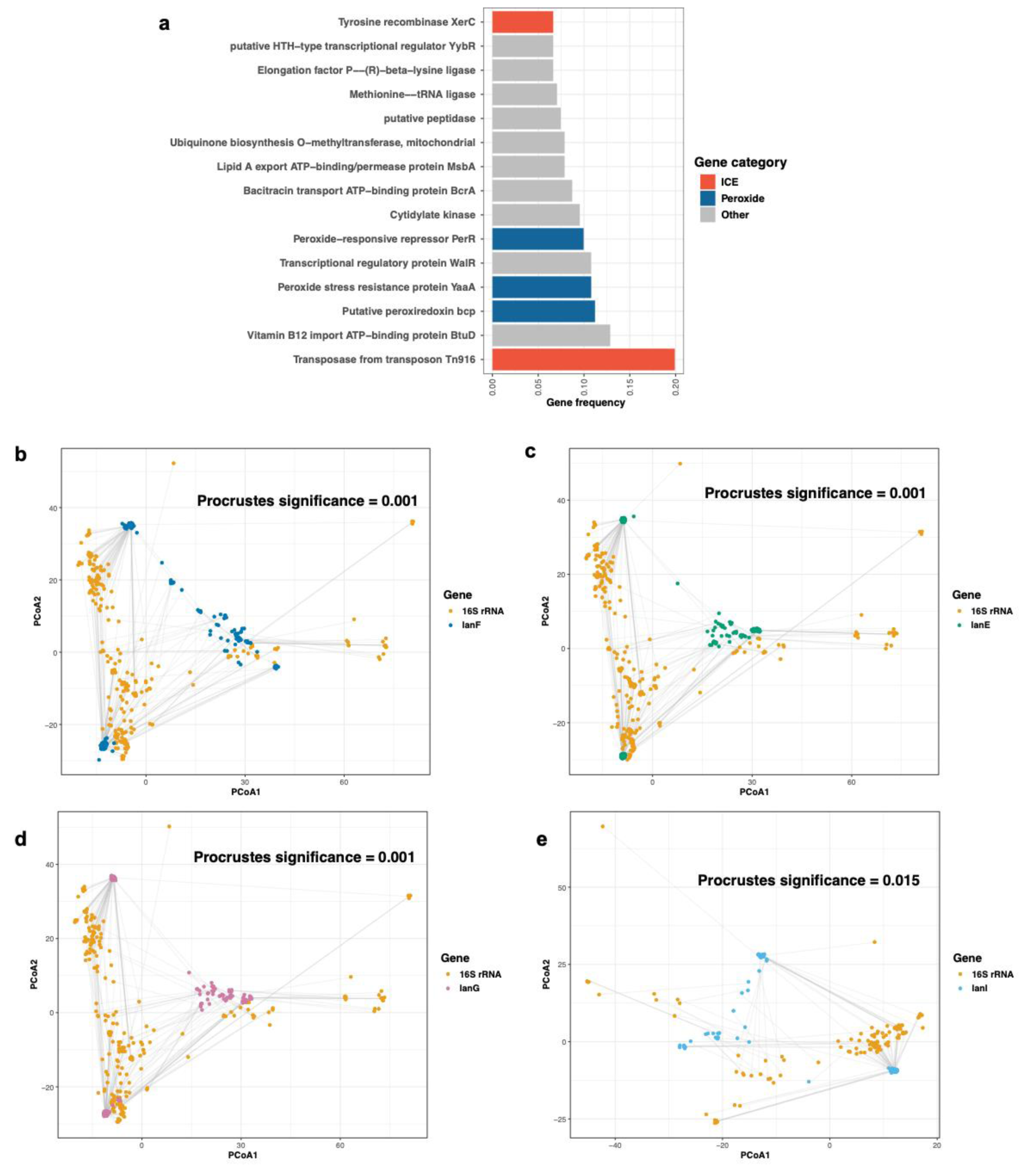
Genomic context of LRS and co-speciation of LRS with host. **(a)** Genomic context of LRS genes. Top 15 most frequent annotations of flanking genes within 10 open reading frames of all LRS in DFI collection are ranked for their occurrence frequency. Genes associated of Integrative Conjugative Elements (ICE), peroxide stress, and others are colored in red, blue and grey, respectively. **(b,c,d,e)** Procrustes superimposition of PCoA plot of 16S rRNA gene (orange) and that of *lanF* (b), *lanE* (c), *lanG* (d) and *lanI* (e), respectively. Lines connect the same genome on two PCoA plots. Significance test of Procrustes analysis was performed with 999 permutations with row and column swapping.

**Extended Data Figure 4.**
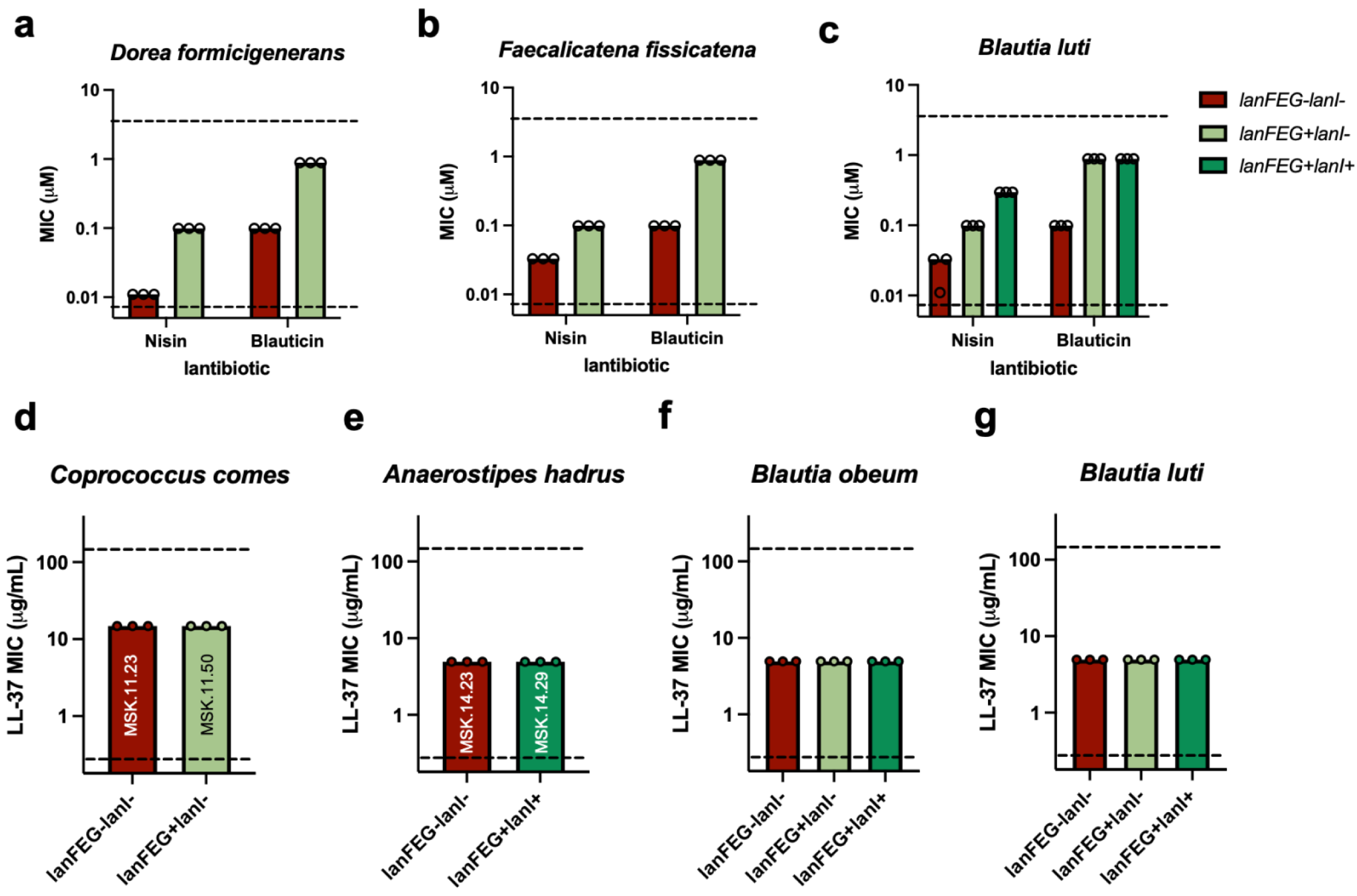
Sensitivity of human gut isolates LRS provides towards lantibiotics and antimicrobial peptides *in vitro*. **(a,b,c)** Minimal Inhibitory Concentration (MIC) of Nisin and Blauticin on *Dorea formicigenerans* (a), *Faecalicatena fissicatena* (b) and *Blautia luti* (c) isolates from human gut. Each circle represents one independent measurement. Horizontal dashed lines indicate upper and lower limit of concentration tested, respectively. Color indicates presence or absence of LRS genes. **(d,e,f,g)** Minimal Inhibitory Concentration (MIC) of human antimicrobial peptide LL-37 on *Coprococcus comes* (d), *Anaerostipes hadrus* (e), *Blautia obeum* (f) and *Blautia luti* (g) isolates from human gut. Each circle represents one independent measurement. Horizontal dashed lines indicate upper and lower limit of concentration tested, respectively. Color indicates presence or absence of LRS genes. Identities of *C. comes* and *A. hadrus* isolates are annotated.

**Extended Data Figure 5.**
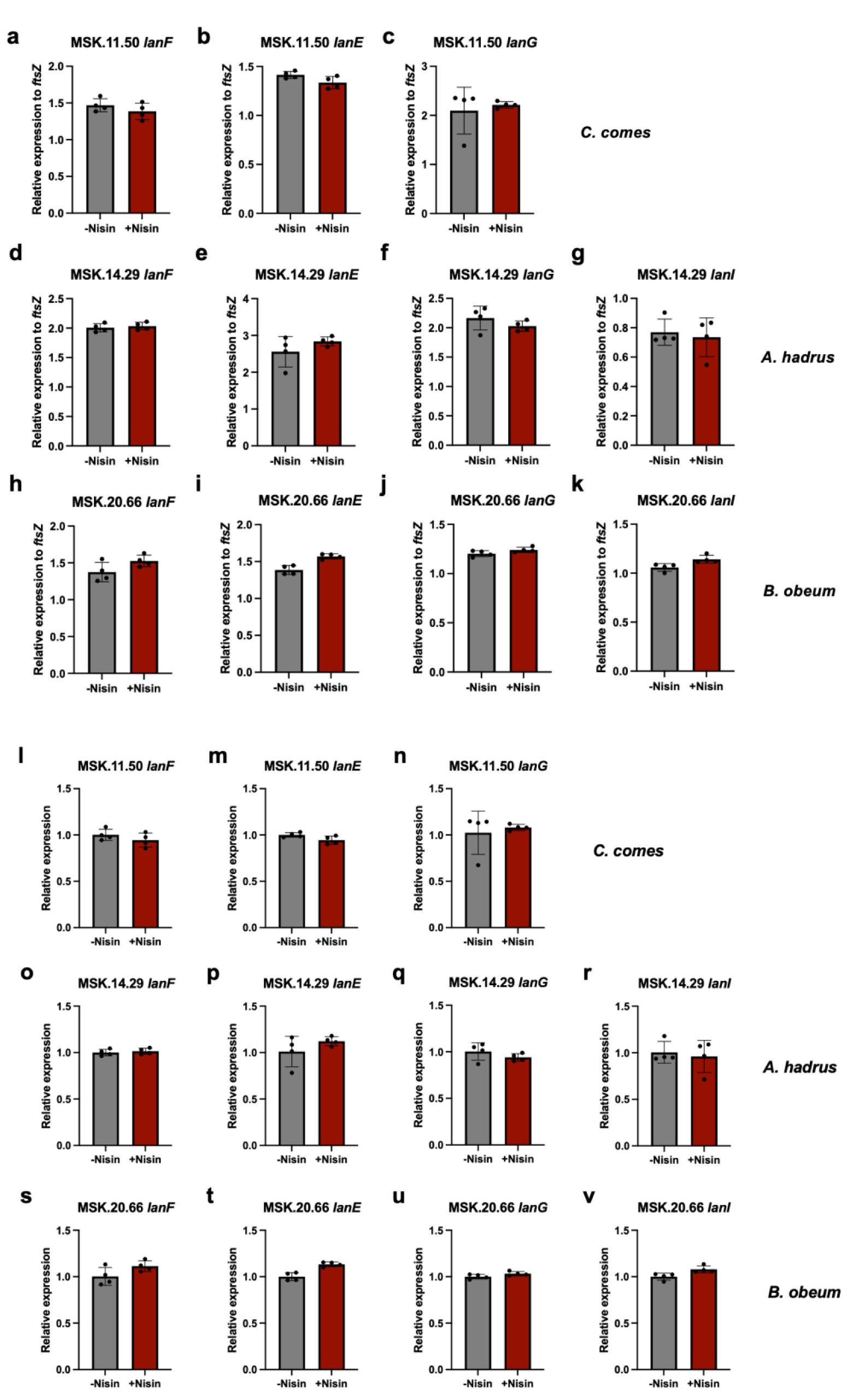
Expression of LRS genes in response to lantibiotic *in vitro*. **(a-k)** Relative expression level of *lanF* (a,d,h), *lanE* (b,e,i), *lanG* (c,f,j) and *lanI* (g,k) in *Coprococcus comes* MSK.11.50 (a,b,c), *Anaerostipes hadrus* MSK.14.29 (d,e,f,g) and *Blautia obeum* MSK.20.66 (h,I,j,k), respectively, in the presence or absence of Nisin, normalized to the expression level of *ftsZ*. Each point represents one biological replicate. Error bars plot standard deviation. Color indicates presence or absence of Nisin in culture. **(l-v)** Relative expression level of *lanF* (l,o,s), *lanE* (m,p,t), *lanG* (n,q,u) and *lanI* (r,v) in *Coprococcus comes* MSK.11.50 (l,m,n), *Anaerostipes hadrus* MSK.14.29 (o,p,q,r) and *Blautia obeum* MSK.20.66 (s,t,u,v), respectively, in the presence or absence of Nisin, normalized to the expression level of respective gene in the absence of Nisin. Each point represents one biological replicate. Error bars plot standard deviation. Color indicates presence or absence of Nisin in culture.

**Extended Data Figure 6.**
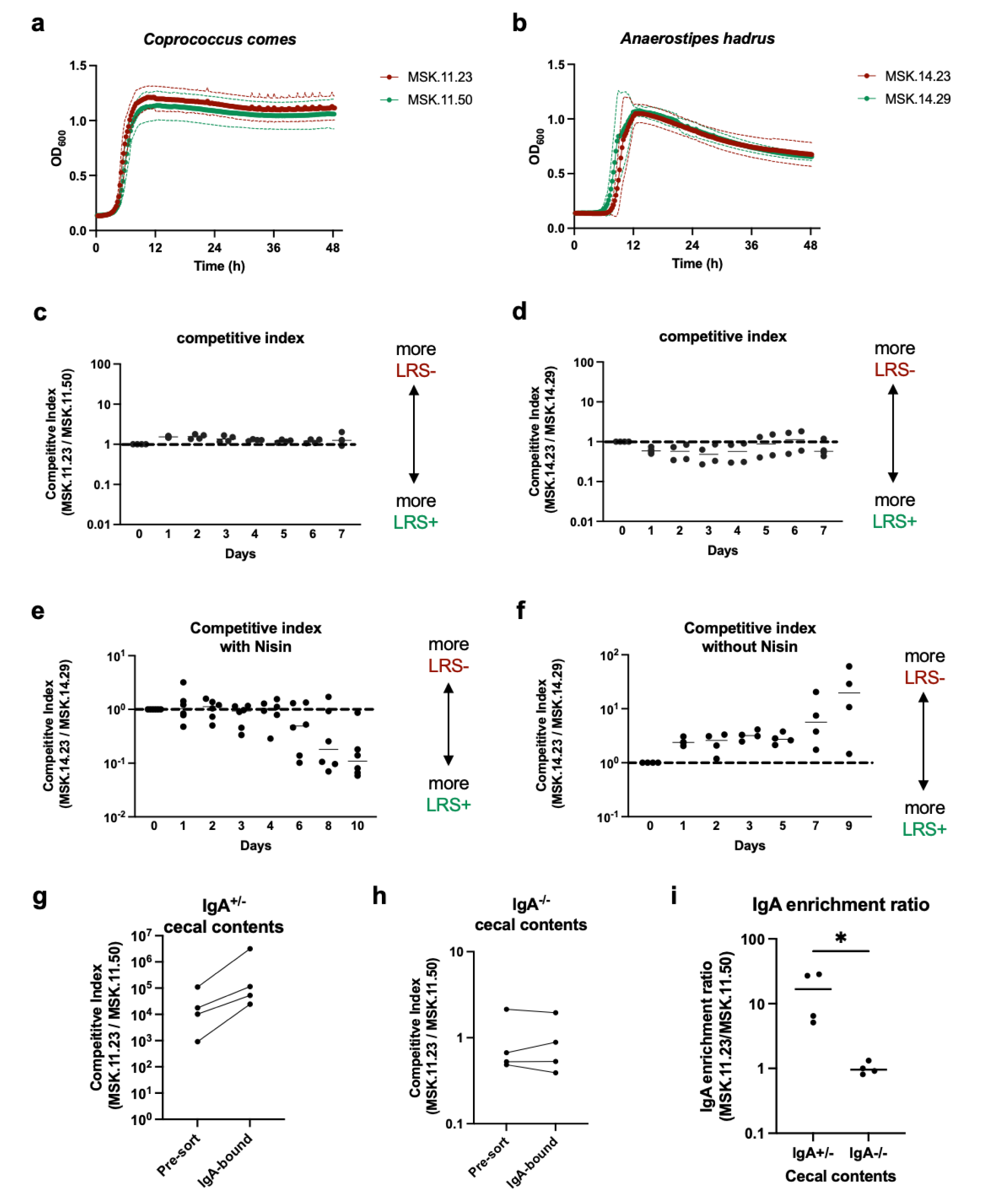
Modulation of fitness dynamics of human gut commensals by LRS. **(a,b)** Growth curves of *Coprococcus comes* (a) and *Anaerostipes hadrus* (b) isolates *in vitro*. Solid points and lines indicate OD_600_ mean, dashed lines indicate standard deviation. LRS-positive and LRS- negative isolates are colored in green and red, respectively. **(c,d)** Competitive index of *C. comes* MSK.11.23 (LRS-negative) over MSK.11.50 (LRS-positive) (c) or that of *A. hadrus* MSK.14.23 (LRS- negative) over MSK.14.29 (LRS-positive) (d) *in vitro*, inoculated with 1:1 ratio of *C. comes* pair or *A. hadrus* pair, and two Bacteroidetes strains on Day 0. Each point represents one biological replicate. Dashed line indicates starting competitive index (1). Solid line plots median at each time point. **(e,f)** Competitive index of *A. hadrus* MSK.14.23 (LRS-negative) over MSK.14.29 (LRS- positive) in gnotobiotic mice with (e) or without Nisin supplementation (f), colonized with 1:1 ratio of *A. hadrus* pair and two Bacteroidetes strains on Day 0. Each point represents one mouse. Dashed line indicates starting competitive index (1). Solid line plots median at each time point. **(g,h)** Competitive index of *C. comes* MSK.11.23 (LRS-negative) over MSK.11.50 (LRS-positive) in cecal contents of IgA^+/-^ (g) or IgA^-/-^ (h) mice before (pre-sort) or after enrichment of IgA-bound fraction (IgA-bound). Each point represents one mouse. Lines connect paired samples. **(i)** IgA enrichment ratio of *C. comes* MSK.11.23 (LRS-negative) over MSK.11.50 (LRS-positive) in cecal contents of IgA^+/-^ or IgA^-/-^ mice, calculated by dividing IgA-bound competitive index by pre-sort competitive index in g and h.

## References

1 Morris, S. & Cerceo, E. Trends, Epidemiology, and Management of Multi-Drug Resistant Gram-Negative Bacterial Infections in the Hospitalized Setting. Antibiotics (Basel) 9 (2020). 10.3390/antibiotics9040196

2 Munk, P. et al. Abundance and diversity of the faecal resistome in slaughter pigs and broilers in nine European countries. Nat Microbiol 3, 898–908 (2018). 10.1038/s41564-018-0192-9

3 Vidovic, N. & Vidovic, S. Antimicrobial Resistance and Food Animals: Influence of Livestock Environment on the Emergence and Dissemination of Antimicrobial Resistance. Antibiotics (Basel) 9 (2020). 10.3390/antibiotics9020052

4 Wang, Y. et al. Comprehensive resistome analysis reveals the prevalence of NDM and MCR-1 in Chinese poultry production. Nat Microbiol 2, 16260 (2017). 10.1038/nmicrobiol.2016.260

5 Mullineaux-Sanders, C., Suez, J., Elinav, E. & Frankel, G. Sieving through gut models of colonization resistance. Nat Microbiol 3, 132–140 (2018). 10.1038/s41564-017-0095-1

6 Le Guern, R. et al. Colonization resistance against multi-drug-resistant bacteria: a narrative review. J Hosp Infect 118, 48–58 (2021). 10.1016/j.jhin.2021.09.001

7 Sorbara, M. T. & Pamer, E. G. Interbacterial mechanisms of colonization resistance and the strategies pathogens use to overcome them. Mucosal Immunol 12, 1–9 (2019). 10.1038/s41385-018-0053-0

8 Chang, P. V. Chemical Mechanisms of Colonization Resistance by the Gut Microbial Metabolome. ACS Chem Biol 15, 1119–1126 (2020). 10.1021/acschembio.9b00813

9 Pike, C. M. & Theriot, C. M. Mechanisms of Colonization Resistance Against Clostridioides difficile. J Infect Dis 223, S194–S200 (2021). 10.1093/infdis/jiaa408

10 Thabit, A. K., Varugehese, C. A. & Levine, A. R. Antibiotic use and duration in association with Clostridioides difficile infection in a tertiary academic medical center: A retrospective case-control study. Anaerobe 59, 126–130 (2019). 10.1016/j.anaerobe.2019.06.016

11 Bell, J. et al. Clostridium difficile Infection Following Spine Surgery: Incidence, Risk Factors, and Association With Preoperative Antibiotic Use. Spine (Phila Pa *1976*) 45, 1572–1579 (2020). 10.1097/BRS.0000000000003627

12 Webb, B. J. et al. Antibiotic Exposure and Risk for Hospital-Associated Clostridioides difficile Infection. Antimicrob Agents Chemother 64 (2020). 10.1128/AAC.02169-19

13 Taur, Y. et al. Intestinal domination and the risk of bacteremia in patients undergoing allogeneic hematopoietic stem cell transplantation. Clin Infect Dis 55, 905–914 (2012). 10.1093/cid/cis580

14 van Nood, E. et al. Duodenal infusion of donor feces for recurrent Clostridium difficile. N Engl J Med 368, 407–415 (2013). 10.1056/NEJMoa1205037

15 Feuerstadt, P. et al. SER-109, an Oral Microbiome Therapy for Recurrent Clostridioides difficile Infection. N Engl J Med 386, 220–229 (2022). 10.1056/NEJMoa2106516

16 Osbelt, L. et al. Klebsiella oxytoca causes colonization resistance against multidrug-resistant K. pneumoniae in the gut via cooperative carbohydrate competition. Cell Host Microbe 29, 1663–1679 e1667 (2021). 10.1016/j.chom.2021.09.003

17 Eberl, C. et al. E. coli enhance colonization resistance against Salmonella Typhimurium by competing for galactitol, a context-dependent limiting carbon source. Cell Host Microbe 29, 1680–1692 e1687 (2021). 10.1016/j.chom.2021.09.004

18 Caballero, S. et al. Cooperating Commensals Restore Colonization Resistance to Vancomycin-Resistant Enterococcus faecium. Cell Host Microbe 21, 592–602 e594 (2017). 10.1016/j.chom.2017.04.002

19 Zhang, Z. J. et al. Structure-Activity Relationship Studies of Novel Gut-derived Lantibiotics Against Human Gut Commensals. bioRxiv, 2023.2009.2015.557961 (2023). 10.1101/2023.09.15.557961

20 Kim, S. G. et al. Microbiota-derived lantibiotic restores resistance against vancomycin-resistant Enterococcus. Nature 572, 665–669 (2019). 10.1038/s41586-019-1501-z

21 Cotter, P. D., Hill, C. & Ross, R. P. Bacteriocins: developing innate immunity for food.Nat Rev Microbiol 3, 777–788 (2005). 10.1038/nrmicro1273

22 Delves-Broughton, J., Blackburn, P., Evans, R. J. & Hugenholtz, J. Applications of the bacteriocin, nisin. Antonie Van Leeuwenhoek 69, 193–202 (1996). 10.1007/BF00399424

23 Bierbaum, G. & Sahl, H. G. Lantibiotics: mode of action, biosynthesis and bioengineering. Curr Pharm Biotechnol 10, 2–18 (2009). 10.2174/138920109787048616

24 Breukink, E. et al. Use of the cell wall precursor lipid II by a pore-forming peptide antibiotic. Science 286, 2361–2364 (1999). 10.1126/science.286.5448.2361

25 Breukink, E. & de Kruijff, B. Lipid II as a target for antibiotics. Nat Rev Drug Discov 5, 321–332 (2006). 10.1038/nrd2004

26 Draper, L. A., Cotter, P. D., Hill, C. & Ross, R. P. Lantibiotic resistance. Microbiol Mol Biol Rev 79, 171–191 (2015). 10.1128/MMBR.00051-14

27 Gharsallaoui, A., Oulahal, N., Joly, C. & Degraeve, P. Nisin as a Food Preservative: Part 1: Physicochemical Properties, Antimicrobial Activity, and Main Uses. Crit Rev Food Sci Nutr 56, 1262–1274 (2016). 10.1080/10408398.2013.763765

28 Blanco-Miguez, A. et al. Extending and improving metagenomic taxonomic profiling with uncharacterized species using MetaPhlAn 4. Nat Biotechnol 41, 1633–1644 (2023). 10.1038/s41587-023-01688-w

29 Sorbara, M. T. et al. Functional and Genomic Variation between Human-Derived Isolates of Lachnospiraceae Reveals Inter- and Intra-Species Diversity. Cell Host Microbe 28, 134–146 e134 (2020). 10.1016/j.chom.2020.05.005

30 Louis, P. & Flint, H. J. Formation of propionate and butyrate by the human colonic microbiota. Environ Microbiol 19, 29–41 (2017). 10.1111/1462-2920.13589

31 Funabashi, M. et al. A metabolic pathway for bile acid dehydroxylation by the gut microbiome. Nature 582, 566–570 (2020). 10.1038/s41586-020-2396-4

32 Sato, Y. et al. Novel bile acid biosynthetic pathways are enriched in the microbiome of centenarians. Nature 599, 458–464 (2021). 10.1038/s41586-021-03832-5

33 Buffie, C. G. et al. Precision microbiome reconstitution restores bile acid mediated resistance to Clostridium difficile. Nature 517, 205–208 (2015). 10.1038/nature13828

34 Foley, M. H. et al. Bile salt hydrolases shape the bile acid landscape and restrict Clostridioides difficile growth in the murine gut. Nat Microbiol 8, 611–628 (2023). 10.1038/s41564-023-01337-7

35 Lloyd-Price, J. et al. Strains, functions and dynamics in the expanded Human Microbiome Project. Nature 550, 61–66 (2017). 10.1038/nature23889

36 Le Chatelier, E. et al. Richness of human gut microbiome correlates with metabolic markers. Nature 500, 541–546 (2013). 10.1038/nature12506

37 Nishijima, S. et al. The gut microbiome of healthy Japanese and its microbial and functional uniqueness. DNA Res 23, 125–133 (2016). 10.1093/dnares/dsw002

38 Smits, S. A. et al. Seasonal cycling in the gut microbiome of the Hadza hunter-gatherers of Tanzania. Science 357, 802–806 (2017). 10.1126/science.aan4834

39 Conteville, L. C., Oliveira-Ferreira, J. & Vicente, A. C. P. Gut Microbiome Biomarkers and Functional Diversity Within an Amazonian Semi-Nomadic Hunter-Gatherer Group. Front Microbiol 10, 1743 (2019). 10.3389/fmicb.2019.01743

40 Hutchinson, M. C., Cagua, E. F., Balbuena, J. A., Stouffer, D. B. & Poisot, T. paco: implementing Procrustean Approach to Cophylogeny in R. Methods in Ecology and Evolution 8, 932–940 (2017). 10.1111/2041-210X.12736

41 Lee, H., Wu, C., Desormeaux, E. K., Sarksian, R. & van der Donk, W. A. Improved production of class I lanthipeptides in Escherichia coli. Chem Sci 14, 2537–2546 (2023). 10.1039/d2sc06597e

42 Vikeli, E. et al. In Situ Activation and Heterologous Production of a Cryptic Lantibiotic from an African Plant Ant-Derived Saccharopolyspora Species. Appl Environ Microbiol 86 (2020). 10.1128/AEM.01876-19

43 Hatziioanou, D. et al. Discovery of a novel lantibiotic nisin O from Blautia obeum A2-162, isolated from the human gastrointestinal tract. Microbiology (Reading) 163, 1292–1305 (2017). 10.1099/mic.0.000515

44 Kuipers, O. P., Beerthuyzen, M. M., Siezen, R. J. & De Vos, W. M. Characterization of the nisin gene cluster nisABTCIPR of Lactococcus lactis. Requirement of expression of the nisA and nisI genes for development of immunity. Eur J Biochem 216, 281–291 (1993). 10.1111/j.1432-1033.1993.tb18143.x

45 Kuipers, O. P., Beerthuyzen, M. M., de Ruyter, P. G., Luesink, E. J. & de Vos, W. M. Autoregulation of nisin biosynthesis in Lactococcus lactis by signal transduction. J Biol Chem 270, 27299–27304 (1995). 10.1074/jbc.270.45.27299

46 Engelke, G. et al. Regulation of nisin biosynthesis and immunity in Lactococcus lactis 6F3. Appl Environ Microbiol 60, 814–825 (1994). 10.1128/aem.60.3.814-825.1994

47 Overhage, J. et al. Human host defense peptide LL-37 prevents bacterial biofilm formation. Infect Immun 76, 4176–4182 (2008). 10.1128/IAI.00318-08

48 Gong, T., Fu, J., Shi, L., Chen, X. & Zong, X. Antimicrobial Peptides in Gut Health: A Review. Front Nutr 8, 751010 (2021). 10.3389/fnut.2021.751010

49 Salzman, N. H. et al. Enteric defensins are essential regulators of intestinal microbial ecology. Nat Immunol 11, 76–83 (2010). 10.1038/ni.1825

50 Fadlallah, J. et al. Microbial ecology perturbation in human IgA deficiency. Sci Transl Med 10 (2018). 10.1126/scitranslmed.aan1217

51 Donaldson, G. P. et al. Gut microbiota utilize immunoglobulin A for mucosal colonization. Science 360, 795–800 (2018). 10.1126/science.aaq0926

52 Palm, N. W. et al. Immunoglobulin A coating identifies colitogenic bacteria in inflammatory bowel disease. Cell 158, 1000–1010 (2014). 10.1016/j.cell.2014.08.006

53 Shapiro, J. M. et al. Immunoglobulin A Targets a Unique Subset of the Microbiota in Inflammatory Bowel Disease. Cell Host Microbe 29, 83–93 e83 (2021). 10.1016/j.chom.2020.12.003

54 Lynch, J. B. et al. Gut microbiota Turicibacter strains differentially modify bile acids and host lipids. Nat Commun 14, 3669 (2023). 10.1038/s41467-023-39403-7

55 Jiao, N. et al. Gut microbiome may contribute to insulin resistance and systemic inflammation in obese rodents: a meta-analysis. Physiol Genomics 50, 244–254 (2018). 10.1152/physiolgenomics.00114.2017

56 Jones-Hall, Y. L., Kozik, A. & Nakatsu, C. Ablation of tumor necrosis factor is associated with decreased inflammation and alterations of the microbiota in a mouse model of inflammatory bowel disease. PLoS One 10, e0119441 (2015). 10.1371/journal.pone.0119441

57 Ross, B. D. et al. Human gut bacteria contain acquired interbacterial defence systems. Nature 575, 224–228 (2019). 10.1038/s41586-019-1708-z

58 Wong, F. et al. Reactive metabolic byproducts contribute to antibiotic lethality under anaerobic conditions. Mol Cell 82, 3499–3512 e3410 (2022). 10.1016/j.molcel.2022.07.009

59 Dwyer, D. J. et al. Antibiotics induce redox-related physiological alterations as part of their lethality. Proc Natl Acad Sci U S A 111, E2100–2109 (2014). 10.1073/pnas.1401876111

60 Zhang, Z. J., Lehmann, C. J., Cole, C. G. & Pamer, E. G. Translating Microbiome Research From and To the Clinic. Annu Rev Microbiol 76, 435–460 (2022). 10.1146/annurev-micro-041020-022206

61 Oliveira, R. A. & Pamer, E. G. Assembling symbiotic bacterial species into live therapeutic consortia that reconstitute microbiome functions. Cell Host Microbe 31, 472–484 (2023). 10.1016/j.chom.2023.03.002

62 Sorbara, M. T. & Pamer, E. G. Microbiome-based therapeutics. Nat Rev Microbiol 20, 365–380 (2022). 10.1038/s41579-021-00667-9

