## Supplementary material for "Exposure and resistance to lantibiotics impact microbiota composition and function": Methods

### Methods for

**Authors and Affiliations**

**Duchossois Family Institute, University of Chicago, 900 E. 57th St, Chicago, IL 60637, USA**
Zhenrun J. Zhang, Huaiying Lin, Cody Cole, Fidel Haro, Emma McSpadden, Eric G. Pamer

**Department of Medicine, Section of Infectious Diseases & Global Health, University of Chicago Medicine, 5841 South Maryland Ave, Chicago, IL 60637, USA**
Eric G. Pamer

**Department of Microbiology, Biological Sciences Division, University of Chicago, 5841 South Maryland Ave, Chicago, IL 60637, USA**
Zhenrun J. Zhang, Cody Cole, Eric G. Pamer

**Department of Chemistry, University of Illinois Urbana-Champaign, IL 61801, USA**
Chunyu Wu, Wilfred A. van der Donk

**Howard Hughes Medical Institute, University of Illinois Urbana-Champaign, IL 61801, USA**
Wilfred A. van der Donk

### Methods

#### Animals

Specific Pathogen Free (SPF) C57BL/6J mice were purchased from Jackson Labs Room AX8 and were maintained under specific pathogen-free conditions at the University of Chicago. Germ Free (GF) C57BL/6J mice were maintained under germ-free conditions at the University of Chicago. GF $IgA^{+/-}$ and $IgA^{+/-}$ mice were kindly gifted by Z. Earley and A. Bendelac lab and were maintained under germ-free conditions at the University of Chicago.

#### SCFA and bile acid profiling of mice fecal samples

SPF C57BL/6J female mice of about 8 weeks old were purchased from Jackson Labs Room AX8. Mice were single-housed and maintained with double-distilled water or water with 10 mg/L Nisin (400 mg/L of 2.5% Nisin, Sigma) for 14 days. Fecal samples were collected on first day (day 0) or 14 days after (day 14) before euthanasia.

To measure concentrations of SCFA, extraction solvent (80% methanol spiked with internal standards and stored at -80°C) was added to pre-weighed fecal samples at a ratio of 100 mg of material/mL of extraction solvent in beadruptor tubes (Fisherbrand; 15-340-154). Samples were homogenized at 4°C on a Bead Mill 24 Homogenizer (Fisher; 15-340-163), set at 1.6 m/s with 6 thirty-second cycles, 5 seconds off per cycle. Samples were then centrifuged at -10°C, 20,000 x g for 15 min and the supernatant was used for subsequent metabolomic analysis. Metabolites were derivatized as described by Haak *et al.* with the following modifications.[1] The metabolite extract (100 µL) was added to 100 µL of 100 mM borate buffer (pH 10) (Thermo Fisher, 28341), 400 µL of 100 mM pentafluorobenzyl bromide (Millipore Sigma; 90257) in Acetonitrile (Fisher; A955-4), and 400 µL of n-hexane (Acros Organics; 160780010) in a capped mass spec autosampler vial (Microliter; 09-1200). Samples were heated in a thermomixer C (Eppendorf) to 65°C for 1 hour while shaking at 1300 rpm. After cooling to room temperature, samples were centrifuged at 4°C, 2000 x g for 5 min, allowing phase separation. The hexanes phase (100 µL) (top layer) was transferred to an autosampler vial containing a glass insert and the vial was sealed. Another 100 µL of the hexanes phase was diluted with 900 µL of n-hexane in an autosampler vial. Concentrated and dilute samples were analyzed using a GC-MS (Agilent 7890A GC system, Agilent 5975C MS detector) operating in negative chemical ionization mode, using a HP-5MSUI column (30 m x 0.25 mm, 0.25 µm; Agilent Technologies 19091S-433UI), methane as the reagent gas (99.999% pure) and 1 µL split injection (1:10 split ratio). Oven ramp parameters were as follows: 1 min hold at 60°C, 25°C per min up to 300°C with a 2.5 min hold at 300°C. Inlet temperature was 280°C and transfer line was 310°C. For quantitative metabolomics of SCFAs, a 10-point calibration curve was prepared with acetate (100 mM), propionate (25 mM), butyrate (12.5 mM), and succinate (50 mM), with 9 subsequent 2x serial dilutions. Data analysis was performed using MassHunter Quantitative Analysis software (version B.10, Agilent Technologies) and confirmed by comparison to authentic standards. Normalized peak areas were calculated by dividing raw peak areas of targeted analytes by averaged raw peak areas of internal standards, which were then converted to quantitative values in mM concentration.

To measure concentrations of bile acids, mice fecal samples were extracted as described above and were analyzed using LC-MS. The metabolite extract (75 µL) was added to prelabeled mass spectrometry autosampler vials (Microliter; 09-1200) and dried down completely under a nitrogen stream at 30 L/min (top) 1 L/min (bottom) at 30°C (Biotage SPE Dry 96 Dual; 3579M). Samples were resuspended in 50:50 Water:Methanol (750 µL). Vials were added to a thermomixer C (Eppendorf) to resuspend analytes at 4°C, 1000 rpm for 15 min with an infinite hold at 4°C. Samples were then transferred to prelabeled microcentrifuge tubes and centrifuged at 4°C, 20,000 x g for 15 min to remove insoluble debris. The supernatant (700 µL) was transferred to a fresh, prelabeled mass spectrometry autosampler vial. Samples were analyzed on a liquid chromatography system (Agilent 1290 infinity II) coupled to a quadrupole time-of-flight (QTOF) mass spectrometer (Agilent 6546), operating in negative mode, equipped with an Agilent Jet Stream Electrospray Ionization source. The sample (5 µL) was injected onto an XBridge© BEH C18 Column (3.5 µm, 2.1 x 100 mm; Waters Corporation, PN) fitted with an XBridge© BEH C18 guard (Waters Corporation, PN) at 45 oC. Elution started with 72% A (Water, 0.1% formic acid) and 28% B (Acetone, 0.1% formic acid) with a flow rate of 0.4 mL/min for 1 min and linearly increased to 33% B over 5 min, then linearly increased to 65% B over 14 min. Then the flow rate was increased to 0.6 mL/min and B was increased to 98% over 0.5 min and these conditions were held constant for 3.5 min. Finally, re-equilibration at a flow rate of 0.4 mL/min of 28% B was performed for 3 min. The electrospray ionization conditions were set with the capillary voltage at 3.5 kV, nozzle voltage at 2 kV, and detection window set to 100-1700 m/z with continuous infusion of a reference mass (Agilent ESI TOF Biopolymer Analysis Reference Mix) for mass calibration. A ten-point calibration curve was used for quantitation. Data analysis was performed using MassHunter Profinder Analysis software (version B.10, Agilent Technologies) and confirmed by comparison with authentic standards. Normalized peak areas were calculated by dividing raw peak areas of targeted analytes by averaged raw peak areas of internal standards.

#### Fecal sample donors

Human donors were enrolled in a prospective fecal collection protocol. Donor age and sex were not reported. The prospective fecal collection protocol was approved by the institutional review board at Memorial Sloan Kettering Cancer Center and The University of Chicago. All donors provided written and informed consent for IRB-approved biospecimen collection and analysis (protocols 06-107; IRB20-1384). The study was conducted in accordance with the Declaration of Helsinki.

#### Isolation and growth of gut bacterial isolates

Isolation and growth of gut bacterial isolates are similar to previously described.[2] Isolation and growth of commensal bacteria was performed under anaerobic conditions with 3% $H_2$, 5% $CO_2$ and 92% $N_2$ in an anaerobic chamber (Coy Labs). Fresh donor fecal samples were transferred into anaerobic conditions within 1 h of collection. Fecal samples were resuspended in pre-reduced

PBS and plated on Columbia agar with 5% sheep blood (BBL, BD) or brain-heart infusion (BHI – Difco, BD) agar plates in three serial 10-fold dilutions and incubated at 37°C for 48 – 96 h. Isolated colonies were re-streaked for purity onto Columbia blood agar and frozen in pre-reduced 10% glycerol in PBS. For broth cultures, isolates were grown in pre-reduced BHI supplemented with 5g/L yeast extract (Difco, BD) and 0.1% L-cysteine (Sigma) (BHIS).

#### Whole-genome shotgun sequencing

Bacterial genomic DNA was extracted from bacterial pellets of broth culture using QIAamp DNA Mini Kit (QIAGEN) according to manufacturer's manual. The purified DNA was quantified using a Qubit 2.0 fluorometer. 1000ng of each sample was prepared for sequencing using the QIAseq FX DNA Library Kit (QIAGEN). The protocol was carried out according to manufacturer's instructions for a targeted fragment size of 550bp. Sequencing was performed on the MiSeq or NextSeq platform (Illumina) with a paired-end (PE) kit in pools designed to provide 1-3 million PE reads per sample with read length of 250 or 150 bp. Adapters were trimmed off with Trimmomatic [3] with following parameters: the leading and trailing 3 bp of the sequences were trimmed off, quality was controlled by a sliding window of 4, with an average quality of 15. Moreover, any read that was less than 50 bp long after trimming and quality control were discarded. The remaining high-quality reads were assembled into contigs using SPAdes (v.3.14.0).[4]

#### Metagenomic shotgun sequencing

Bacterial DNA was extracted using the QIAamp PowerFecal Pro DNA kit (Qiagen). Prior to extraction, samples were subjected to mechanical disruption using a bead beating method. Briefly, samples were suspended in a bead tube (Qiagen) along with lysis buffer and loaded on a bead mill homogenizer (Fisherbrand). Samples were then centrifuged, and supernatant was resuspended in a reagent that effectively removed inhibitors. DNA was then purified routinely using a spin column filter membrane and quantified using Qubit. Libraries were prepared using 200 ng of genomic DNA using the QIAseq FX DNA library kit (Qiagen). Briefly, DNA was fragmented enzymatically into shorter fragments and desired insert size was achieved by adjusting fragmentation conditions. Fragmented DNA was end repaired and 'A's' were added to the 3'ends to stage inserts for ligation. During ligation step, Illumina compatible Unique Dual Index (UDI) adapters were added to the inserts and prepared library was PCR amplified. Amplified libraries were cleaned up, and QC was performed using Tapestation 4200 (Agilent Technologies). Libraries were sequenced on an Illumina MiSeq or NextSeq 500 platforms to generate 2x250bp or 2x150bp reads, respectively, targeted for 2 to 3.2 million pair-end reads per sample.

#### Metagenomic reads mapping

To quantify the relative abundance of bile salt hydrolases (BSH) and taurine-preferring BSH in metagenomes of mice fecal samples, BSH and taurine-preferring BSH protein sequences were retrieved (Extended Files) [5] and aligned to metagenomic reads of mice fecal samples with or without Nisin exposure using DIAMOND [6]. Aligned reads were filtered with at least 80% identity, at least 80% translated length coverage, and E-value less than 1e-5. Each filtered read were called for the best protein hit according to highest bitscore, highest percentage identity and lowest E-value, in that order. RPKM were tallied with RPKM of each read according to length of its mapped protein sequence and the read count of the metagenomic sample.

To quantify the relative abundance of lantibiotic resistance genes (LRG) and *lanA* in metagenomes of mice fecal samples and in metagenomes of human cohorts, LRG protein sequences were compiled from unique LRG protein sequences from RefSeq and this study (Extended Files). To compile a list of *lanA* protein sequences, non-redundant bacterial protein sequences of RefSeq (release 205) were searched with Hidden Markov Model of Type-A lantibiotic (PF04604.14) using HMMER3 [7], and hits with E-value less than 1e-3 were retrieved (Extended File). Metagenomes of human cohorts were retrieved from National Center for Biotechnology Information projects: PRJDB3601 (Nishijima et al., 2016), PRJNA48479 (Lloyd-Price et al., 2017), PRJEB4336 (Le Chatelier et al., 2013), PRJEB2054 (Qin et al., 2010), PRJNA392180 (Smits et al., 2017), and PRJNA527208 (Conteville et al., 2019). LRG and *lanA* sequences were aligned to metagenomic reads of mice fecal samples with or without Nisin exposure and human metagenomes using DIAMOND [6] and were filtered and quantified using similar workflow described above.

#### Genomic and LRG annotation and LRG context

For individual isolates, the genome assemblies were annotated using Prokka (v. 1.12)[8]. To find and annotate LRG in isolates' genomes, Hidden Markov Model (HMM) of LanF, LanE and LanG proteins were built using seed sequences from RefSeq. HMM of LanI protein was retrieved from Pfam (Spa1_C, PF18218). Annotated genomes were searched for homologs of LanF, LanE, LanG and LanI, respectively, using HMMER3 [7]. For potential hits of LanF, LanE or LanG in the same genome, their genomic locations were checked so that only consecutive *lanFEG* genes in that order were annotated as bona fide LRG. After consolidating LRG in genomes, their neighboring genes within 10 open reading frames were tallied for frequency of annotations, excluding hypothetical proteins.

#### Procrustes analysis

For Procrustes cophylogeny test between LRG and their hosts, pairwise distance matrix of each LRG and host longest 16S rRNA sequences were calculated based on Multiple Sequence Alignment (MSA) by Clustal Omega [9]. Procrustes cophylogeny tests between 16S rRNA gene and each LRG were performed using PACo (v0.4.2)[10], using distance matrix of 16S rRNA sequences as "host" and distance matrix of each LRG as "parasite". PACo test was performed with row and column swapping (method = 'r00') and 1000 permutations (nperm = 1000).

#### Minimal Inhibition Concentration (MIC) assays

All MIC assays were performed under anaerobic condition (3% $H_2$, 5% $CO_2$ and 92% $N_2$). Stationary bacterial culture in BHIS (BHI supplemented with 5g/L yeast extract and 0.1% L-cysteine) were inoculated 1:100 to wells containing 200 uL BHIS in U-bottom 96-well plates.

Lantibiotics and AMPs were dissolved in DPBS, and concentrations were determined by BCA assay (Pierce) according to manufacturer's protocol. Lantibiotic stock solutions were added to wells to reach desired final concentration, with 3-fold serial dilution across rows to create concentration gradients. Bacteria-inoculated wells without lantibiotics served as control. Plates were incubated at 37$^{o}$C for 48 h before taken out of anaerobic chamber for centrifugation. Plates were imaged with iBright 1500 imager (Invitrogen). The lowest concentration of lantibiotics and AMPs that caused no or significantly less bacterial pellets than those without lantibiotics were identified as MIC.

#### Gnotobiotic mice experiments

For gnotobiotic mice experiments that monitor the competition between the pair of LRG-containing and LRG-lacking strains, the pair of strains (*C. comes* MSK.11.23 and MSK.11.50; or *A. hadrus* MSK.14.23 and MSK.14.29) as well as supporting Bacteroidetes strains (*Bacteroides sartorii* CBBP and *Parabacteroides distasonis* CBBP {Caballero, 2017 #19;Kim, 2019 #18}) were individually culture in BHIS for 48 h before $OD_{600}$ measurement. Assuming all culture has a density of 1 x $10^8$ CFU/mL when $OD_{600}$ = 1, 1 x $10^8$ CFU of each bacteria were aliquoted according their $OD_{600}$ value before 10,000 x g centrifugation for 5 min. Supernatants were discarded and bacterial pellets were resuspended and mixed in 1 mL pre-reduced DPBS. Bacterial consortium suspension was stored in airtight O-ring centrifuge tubes on ice and were opened right before gavage. Germ-free (GF) C57BL/6J WT, $IgA^{+/-}$ or $IgA^{+/-}$ mice, male and female aged about 8 weeks, were transferred from GF mice facility to clean hood before oral gavage of 200 uL of bacterial consortium suspension (2 x $10^7$ CFU of each bacteria). Mice were single housed in hermetically sealed cages and their fecal samples were collected and stored in -80$^{o}$C freezer according to day schedule. For experiments with Nisin supplementation, one day after gavage of bacterial consortium suspension, drinking water was switched to water containing 10 mg/L Nisin (400 mg/L 2.5% Nisin, Sigma).

#### Quantitative Reverse-Transcription PCR and Competitive Index

#### References


1 Haak, B. W. *et al.* Impact of gut colonization with butyrate-producing microbiota on respiratory viral infection following allo-HCT. *Blood* **131**, 2978-2986 (2018). https://doi.org/10.1182/blood-2018-01-828996

2 Sorbara, M. T. *et al.* Functional and Genomic Variation between Human-Derived Isolates of Lachnospiraceae Reveals Inter- and Intra-Species Diversity. *Cell Host Microbe* **28**, 134-146 e134 (2020). https://doi.org/10.1016/j.chom.2020.05.005

3 Bolger, A. M., Lohse, M. & Usadel, B. Trimmomatic: a flexible trimmer for Illumina sequence data. *Bioinformatics* **30**, 2114-2120 (2014). https://doi.org/10.1093/bioinformatics/btu170

4 Bankevich, A. *et al.* SPAdes: a new genome assembly algorithm and its applications to single-cell sequencing. *J Comput Biol* **19**, 455-477 (2012). https://doi.org/10.1089/cmb.2012.0021

5 Foley, M. H. *et al.* Bile salt hydrolases shape the bile acid landscape and restrict Clostridioides difficile growth in the murine gut. *Nat Microbiol* **8**, 611-628 (2023). https://doi.org/10.1038/s41564-023-01337-7

6 Buchfink, B., Reuter, K. & Drost, H. G. Sensitive protein alignments at tree-of-life scale using DIAMOND. *Nat Methods* **18**, 366-368 (2021). https://doi.org/10.1038/s41592-021-01101-x

7 Eddy, S. R. Accelerated Profile HMM Searches. *PLoS Comput Biol* **7**, e1002195 (2011). https://doi.org/10.1371/journal.pcbi.1002195

8 Seemann, T. Prokka: rapid prokaryotic genome annotation. *Bioinformatics* **30**, 2068-2069 (2014). https://doi.org/10.1093/bioinformatics/btu153

9 Sievers, F. & Higgins, D. G. Clustal Omega for making accurate alignments of many protein sequences. *Protein Sci* **27**, 135-145 (2018). https://doi.org/10.1002/pro.3290

10 Hutchinson, M. C., Cagua, E. F., Balbuena, J. A., Stouffer, D. B. & Poisot, T. paco: implementing Procrustean Approach to Cophylogeny in R. *Methods in Ecology and Evolution* **8**, 932-940 (2017). https://doi.org/https://doi.org/10.1111/2041-210X.12736

11 Eren, A. M. *et al.* Community-led, integrated, reproducible multi-omics with anvi'o. *Nat Microbiol* **6**, 3-6 (2021). https://doi.org/10.1038/s41564-020-00834-3

12 McInnes, L., Healy, J. & Melville, J. Umap: Uniform manifold approximation and projection for dimension reduction. *arXiv preprint arXiv:1802.03426* (2018).

13 Yin, Y. *et al.* dbCAN: a web resource for automated carbohydrate-active enzyme annotation. *Nucleic Acids Res* **40**, W445-451 (2012). https://doi.org/10.1093/nar/gks479

14 Stewart, R. D., Auffret, M. D., Roehe, R. & Watson, M. Open prediction of polysaccharide utilisation loci (PUL) in 5414 public <em>Bacteroidetes</em> genomes using PULpy. *bioRxiv*, 421024 (2018). https://doi.org/10.1101/421024

15 Garcia-Bayona, L., Coyne, M. J. & Comstock, L. E. Mobile Type VI secretion system loci of the gut Bacteroidales display extensive intra-ecosystem transfer, multi-species spread and geographical clustering. *PLoS Genet* **17**, e1009541 (2021). https://doi.org/10.1371/journal.pgen.1009541

16 Coyne, M. J., Roelofs, K. G. & Comstock, L. E. Type VI secretion systems of human gut Bacteroidales segregate into three genetic architectures, two of which are contained on mobile genetic elements. *BMC Genomics* **17**, 58 (2016). https://doi.org/10.1186/s12864-016-2377-z

17 Darzi, Y., Falony, G., Vieira-Silva, S. & Raes, J. Towards biome-specific analysis of meta-omics data. *ISME J* **10**, 1025-1028 (2016). https://doi.org/10.1038/ismej.2015.188

18 Vieira-Silva, S. *et al.* Species-function relationships shape ecological properties of the human gut microbiome. *Nat Microbiol* **1**, 16088 (2016). https://doi.org/10.1038/nmicrobiol.2016.88
